# Structural basis for a filamentous morpheein model of human cystathionine beta-synthase

**DOI:** 10.1101/2025.09.24.678195

**Authors:** Inayathulla Mohammed, Ela Mijatovic, Thilo M Philipp, Lucia Janickova, Kelly Ascencao, Francisco J Asturias, Csaba Szabo, Henning Stahlberg, Tomas Majtan

## Abstract

Human cystathionine beta-synthase (CBS) is a vital enzyme that regulates sulfur amino acid metabolism, hydrogen sulfide production, and cellular redox balance. Using a multidisciplinary approach, we demonstrate that CBS functions as a filamentous morpheein, with its stability, turnover, and activity governed by dynamic quaternary structural transitions. Three distinct filamentous assemblies were resolved by cryo-EM and are mediated by the oligomerization loop (residues 516–525): (i) ligand-free *trans*-dimers that form *trans*-basal filaments with basal stability and activity, (ii) adenosylornithine-bound *cis*-dimers that assemble into stabilized *cis*-basal filaments and (iii) S-adenosylmethionine-bound *allo*-dimers, which, together with *cis*-dimers, form highly stable, *allo*-activated stacked filaments. These reversible filamentous assemblies redefine CBS biology by integrating oligomerization and allosteric regulation within a morpheein framework. These findings provide a transformative perspective on CBS function and open new avenues for pharmacological targeting of dysregulated CBS in various diseases including homocystinuria, cancer, and Down syndrome.

## Introduction

Human cystathionine beta-synthase (CBS) is an essential enzyme in human metabolism, orchestrating the biogenesis of small redox-active molecules such as cysteine, glutathione, and hydrogen sulfide, which regulate cellular health and signaling [1]. CBS is a complex heme-containing pyridoxal-5’-phosphate-dependent enzyme allosterically activated by S-adenosylmethionine (SAM) (**Figure 1A**). SAM binds to the regulatory domain, which also plays a crucial role in CBS oligomerization and kinetic stabilization [2–6]. Previous study showed that CBS is a filamentous protein [7]; however, the molecular mechanisms controlling structural rearrangements and its function remained unclear. Here we show that CBS operates as a filamentous morpheein, with its stability, turnover, and activity governed by dynamic, reversible transitions between three distinct filament types driven by the character of an allosteric ligand: (i) ligand-free *trans*-basal filaments with basal stability and activity, (ii) non-activating ligand adenosylornithine-bound *cis*-basal filaments with increased stability and basal activity and (iii) SAM-bound *allo*-activated stacked filaments with high stability and activity. These findings reveal a novel regulatory framework, integrating oligomerization and allosteric control, that redefines our understanding of the CBS biology. As dysregulation of CBS stability and activity contributes to homocystinuria, Down syndrome and cancer [2, 8, 9], this discovery advances the field by clarifying how structural dynamics modulate its function and consequently opens new avenues for developing targeted pharmacological therapies.

**Figure 1.**
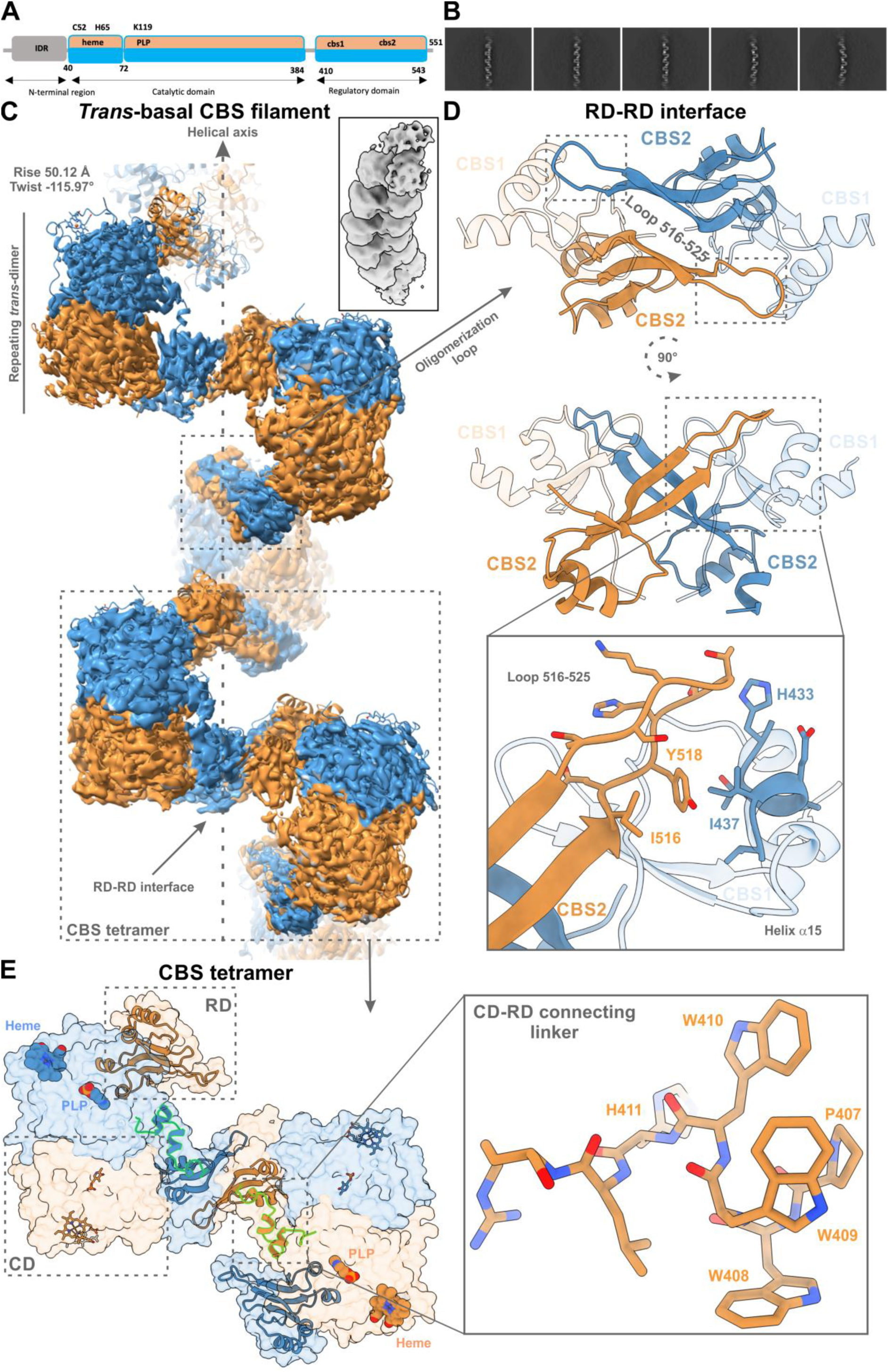
Architecture and inter-domain interfaces of the human CBS *trans*-basal filament. **A** – Schematic representation of human CBS domain organization. **B** – Representative 2D class averages of human CBS filaments in the absence of allosteric ligands. The filaments exhibit variable lengths, reflecting their dynamic and heterogenous nature. **C** – Helical 3D reconstruction of the *trans*-basal CBS filament resolved to a global resolution of 3.0 Å. The inset shows an oblique view of the filament highlighting the helical geometry with a twist of –116° and axial rise of 50 Å per asymmetric unit. The repeating unit is a *trans*-dimer composed of two protomers (blue and orange), arranged with D1 symmetry orthogonal to the helical axis. Oligomerization is mediated by interactions between the regulatory domains (RDs) of *trans*-dimers, interconnected through the oligomerization loop. **D** – Atomic model detailing the interface between RDs of two dimers in the repeating unit. The oligomerization loop (residues 516–525) is crucial for CBS filamentation and hydrophobic contacts between residues Y518 and I516 from the CBS-2 oligomerization loop and residues I437 and H433 of CBS-1 helix α15 further stabilize interaction between CBS dimers (inset). **E** – Space-filled cryo-EM density representation and atomic model illustrating how *trans*-dimers organize into tetrameric assemblies (dimers of dimers) via head-to-tail interactions. The inset provides a close-up view of the linker region connecting the catalytic (CD) and regulatory (RD) domains (residues 384–414). A cluster of tryptophan residues, situated between H411 and P407, are positioned to facilitate allosteric signaling between the CD and RD, highlighting their potential involvement in conformational transitions and allosteric activation of CBS.

## Results

### Human CBS is a filamentous enzyme

Recombinantly expressed, purified human full-length CBS in the absence of SAM (i.e. the basal state) formed ordered filaments with variable length (**Figure 1B**). The cryo-EM helical analysis yielded a reconstruction at a resolution of 3.0 Å, revealing a helical architecture with a twist of -116° and a rise of 50 Å per subunit (**Figure 1C**). The fundamental repeating unit of the filament is a *trans*-CBS dimer, i.e. a dimer, in which the RDs are domain-swapped relative to their respective CDs. This conformation resembles that of a dimeric CBSΔ516-525 variant previously solved by X-ray crystallography in the absence of SAM [10]. A crucial structural feature facilitating filamentous assembly is the oligomerization loop spanning residues 516-525 (**Figure 1D**). This loop mediates inter-dimer interactions resulting in the formation of higher-order tetrameric units (**Figure 1E**). These tetramers act as modular building blocks that extend to form filaments composed of up to 20 such units (**Supplementary figure 2**). The head-to-tail arrangement of dimers along the central axis, stabilized by the 516–525 loop, defines the oligomeric state referred here to as the *trans*-basal CBS assembly (**Figure 1C**). Two key interfaces between RDs contribute to the stabilization of *trans*-basal CBS filament. The first involves interactions between the CBS1 and CBS2 motifs of adjacent subunits, bridged by the 516-525 loop and α-helix 15 (**Figure 1D**). Additional hydrogen bonds and main-chain contacts between β-strands 12 and 8 further strengthen this interface. Within this context, residue Y518 is positioned into a hydrophobic pocket formed by residues within α-helix 15. The second interface consists of hydrophobic packing interactions between residues of adjacent CBS2 motifs, specifically involving L419, L492, M529 and F531. These contacts induce a reorientation of A421 and P422 relative to their positions in the crystal structure of CBSΔ516–525 [10]. Altogether, these inter-dimer interactions, mediated by the oligomerization loop 516-525, are critical for the assembly and stabilization of the *trans*-basal CBS filament under ligand-free conditions.

### The oligomeric status of CBS determines its cellular turnover

Filamentous organization, a feature common among certain metabolic enzymes, has been shown to provide functional advantages, such as rapid deployment, catalytic enhancement or structural stabilization [11]. Given that both CBS WT and CBSΔ516-525 exhibit comparable catalytic activities in the absence and presence of SAM [10], we hypothesized that filamentation may contribute to CBS stabilization in a cellular milieu. To test this hypothesis, we utilized a non-radioactive pulse-labeling method to assess cellular turnover of two annotated CBS isoforms along with two engineered constructs [12]. HEK293 cells lacking endogenous CBS, achieved via CRISPR-Cas9-mediated knockout (**Figure 2A**), were stably transfected to express full-length CBS WT (isoform 1, ISO1), CBS isoform 2 containing additional 15 residues replacing Y518 encoded by an alternative exon 15 (ISO2), CBSΔ516-525 and CBS45 (**Figure 2B**). Notably, ISO1 predominantly formed tetramers and higher-order oligomers, whereas ISO2, CBSΔ516-525 and CBS45 primarily existed as dimers (**Figure 2C**). This observation highlights the critical role of residue Y518, which fits into a hydrophobic pocket within α-helix 15 of the adjacent subunit, in promoting CBS filamentation (**Figure 1D**). All constructs retained catalytic activity and responded to SAM except for CBS45, which does not bind SAM due to the lack of the RD and is constitutively activated (**Figure 2D-E**). More importantly, full-length oligomeric CBS WT (ISO1) showed a cellular half-life of 13.3 h, which was approximately twice that of ISO2 and CBSΔ516-525 (6.6 h and 6.4 h, respectively) and nearly five times longer than the 2.7-hour half-life of CBS45 (**Figure 2F**). These findings suggest that the tetramer observed on native PAGE corresponds to a functional assembly unit (a dimer of dimers) required for filamentation, while the predominant dimer correlated with impaired filamentation. Confocal microscopy analysis further supported these findings. Tetrameric/filamentous CBS WT (ISO1) exhibited concentrated fluorescence near the nucleus, whereas dimeric forms (ISO2, CBSΔ516-525, CBS45) showed a diffuse cytoplasmic distribution (**Figure 2G**). Collectively, these results indicate that CBS tetramerization (and, by extension, filamentation) not only reduces protein turnover but also influences subcellular localization, thereby enhancing the enzyme’s stability and function in the cell.

**Figure 2.**
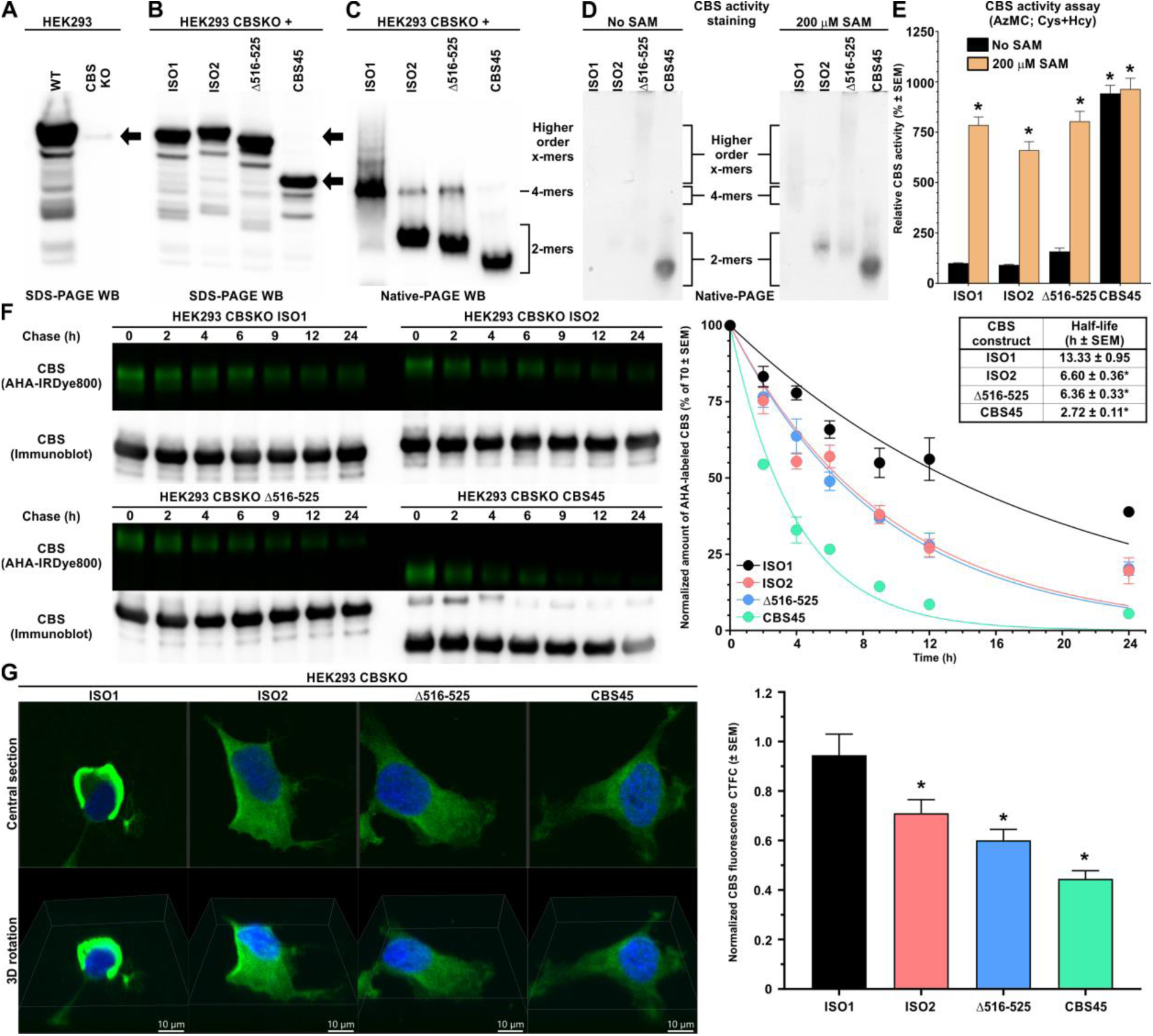
Cellular turnover of CBS is determined by its oligomeric status. (**A**) SDS-PAGE Western blot confirming the absence of endogenous CBS in HEK293 CBSKO cells. (**B**) SDS-PAGE and (**C**) native PAGE Western blots of cell lysates showing expression and oligomeric status, respectively, of the studied CBS constructs. (**D**) In-gel CBS activity staining and (**E**) AzMC-based fluorescence assay in cell lysates in the absence or presence of 200 µM SAM. (**F**) Determination of cellular turnover of the studied CBS constructs showing representative gels and blots (left) used for signal quantification, normalization, plotting of the CBS decay (right) and calculation of CBS half-life (inset). (**G**) Confocal imaging of the studied CBS constructs showing their cellular compartmentalization (green = CBS, blue = nucleus; left) and quantification of the normalized CBS signal (right).

### Substrate binding reveals catalytic rearrangements of the active site

To gain mechanistic insight into CBS catalysis, we determined cryo-EM structures of the enzyme in both the absence and presence of its substrate, L-serine (**Figure 3**). In its absence, the catalytic pocket showed a covalent internal aldimine intermediate formed between the PLP and residue K119 (CBS-Lys-PLP; **Figure 3A**). Upon the addition of serine, we captured two additional structural states. In the first one, serine was covalently bound to PLP forming the external aldimine intermediate (CBS PLP-Ser; **Figure 3B**). The second state showed the elimination of water from the PLP-Ser adduct resulting in the formation of the PLP-aminoacrylate intermediate (CBS PLP-AA; **Figure 3C**). Formation of the PLP-AA intermediate represents a key transition in the catalytic cycle, as it primes CBS to accept the second substrate. The reaction proceeds through a β-replacement or α,β-elimination mechanism using the canonical substrate homocysteine and alternative H_2_S, respectively [13, 14]. Transition from internal aldimine to the PLP-AA intermediate triggered a compaction of the active site, which was reflected in a global contraction of the catalytic pocket and coordinated repositioning of the residues involved in catalysis. This local active site remodeling coincided with an observable movement along the filamentous assembly of CBS. Specifically, subtle rocking-like motions centered around the CDs, transmitted through the flanking RDs along the filamentous axis, suggest a dynamic “breathing” behavior of the *trans*-basal CBS filament (**Supplementary Movies 1&2**). This conformational plasticity may represent an intrinsic feature of the enzyme’s catalytic cycle, potentially coupling local chemical events to long-range structural shifts across the filament.

**Figure 3.**
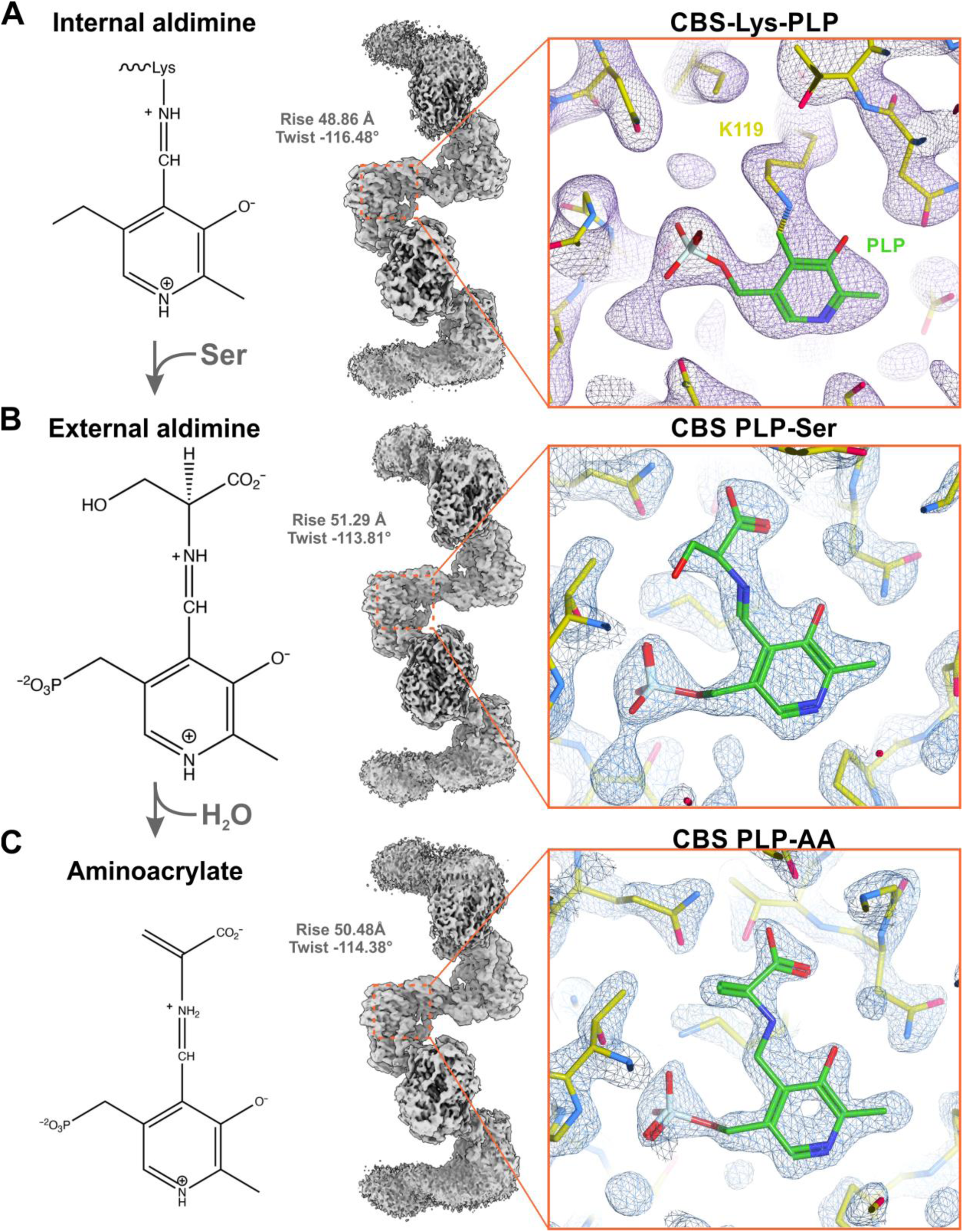
Substrate binding induces active site remodeling and catalytic intermediate formation in CBS. Transition from the substrate-free internal aldimine to the PLP-aminoacrylate (PLP-AA) intermediate results in a marked compaction of the catalytic pocket, accompanied by precise, coordinated rearrangements of the surrounding active site residues. Human CBS was successfully trapped in two distinct intermediate states and resolved at high resolutions of 2.1 Å and 2.3 Å, enabling detailed interpretation of the associated stereochemical transformations. **A** – Cryo-EM structure of CBS in the absence of substrate, revealing the catalytic pocket occupied by the internal aldimine formed between the PLP cofactor and residue K119. This represents the resting, unprimed state of the enzyme (CBS-Lys-PLP). **B** – Structure of serine-bound CBS showing formation of the PLP-serine external aldimine intermediate. Serine is covalently linked to PLP, marking the first step of substrate engagement in the CBS catalytic cycle (CBS PLP-Ser). **C** – Structure of CBS PLP-aminoacrylate intermediate (CBS PLP-AA) formed after elimination of water from the PLP-Ser adduct. This state represents the key intermediate making CBS active site primed for accepting the second substrate and follow either the β-replacement or α,β-elimination.

### Non-activating allosteric ligand induces a distinct CBS filamentous state

To investigate how binding of allosteric ligands affects the morphology of the CBS filament, first we examined the impact of adenosylornithine (sinefungin, SAO), a CBS non-activating SAM analog (**Figure 4**). Previous crystallographic study using CBSΔ516-525 E201S variant demonstrated that SAM binds to the S2 site within the RD with a 1:1 stoichiometry per monomer [3]. This interaction promoted a substantial rearrangement of the RDs, into antiparallel, head-to-tail CBS modules. The structural reorganization, facilitated by a flexible interdomain linker (**Figure 1E**), resulted in the spatial separation of the RDs from the CDs. However, it remained unclear whether the previously studied SAM analogs, such as S-adenosylhomocysteine (SAH) and SAO (**Figure 4A**), could elicit similar structural changes [5]. The presence of SAH and SAO did not activate CBS (**Figure 4B**) suggesting that mere occupation of the binding site may not be sufficient for the activation and that specific conformational rearrangements are essential to achieve that. Thus, we initially anticipated that SAO binding would not induce notable quaternary rearrangements. Contrary to this expectation, cryo-EM analysis of CBS WT in the presence of SAO revealed a novel filament morphology, distinct from the *trans*-basal state (**Figure 4C**). Due to filament flexibility, localized 3D refinement was necessary to obtain a composite global reconstruction at 4.0 Å resolution. The structure revealed a central helical core formed by antiparallel RD dimers flanked on either side by dimers of CDs. This repeating unit, distinct from the trans-dimer was defined as the *cis*-CBS dimer (**Figure 4D–F**). Interestingly, this architecture closely resembled that of the constitutively activated CBSΔ516–525 E201S in the SAM-bound state [3], which previously led to the hypothesis that such a configuration may represent the activated form of full-length CBS [7]. Our current data, however, indicate that SAO binds to the same S2 site as SAM (**Figure 4F-H**), but fails to activate the enzyme (**Figure 4B**) [5, 15]. Thus, despite the structural rearrangement, CBS remained catalytically unaltered in the presence of SAO. Accordingly, we designated this assembly as the *cis*-basal CBS filament. The *cis*-basal CBS filament exhibited a straight geometry with an overall right-handed helical twist of -173° and an axial rise of 49 Å. While the CDs retain minimal contacts with the central stalk consisting of the RDs, consistent with the previous activation models [3, 7], our structure revealed an absence of any functional coupling. Specifically, the lack of interactions between RDs and the flexible loops located near the catalytic entrance indicates that the additional structural determinants, likely specific to SAM-induced assemblies, are essential for CBS allosteric activation. Additionally, we also resolved a medium-resolution map capturing a transient intermediate during SAO binding (**Supplementary figure 14B**). This structure resembles a transitional state between the ligand-free *trans*-basal and SAO-bound *cis*-basal conformations, representing an intermediate step in the ligand-induced rearrangement.

**Figure 4.**
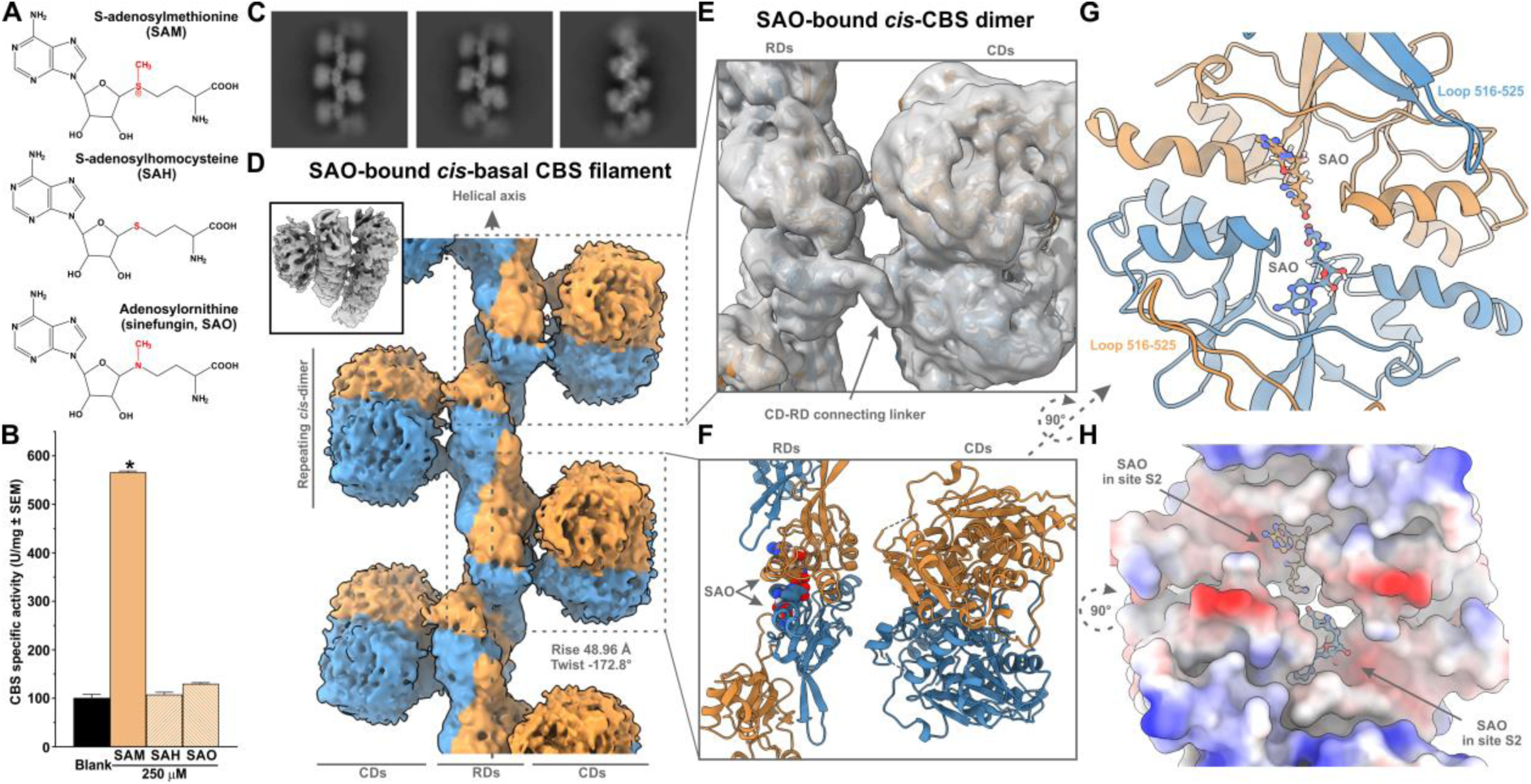
Structural characterization of the *cis*-basal CBS filament induced by the CBS non-activating allosteric ligand adenosylornithine (SAO). A – Chemical structures of SAM, SAH and SAO highlighting the key structural differences (in red). **B** – CBS specific activity in the absence (blank) or presence of 250 µM allosteric ligands demonstrating that SAO does not activate CBS. **C** – Representative 2D class averages of CBS filaments assembled in the presence of SAO, showing a distinct morphology compared to the *trans*-basal filament observed in the absence of allosteric ligands. **D** – Helical cryo-EM reconstruction of the *cis*-basal CBS filament at an overall resolution of 4.0 Å. The inset shows an oblique view of the filament highlighting its right-handed helical geometry with a twist of –173° and axial rise of 49 Å. The repeating unit is a *cis*-dimer composed of two protomers (blue and orange), where their RDs forming disc-shaped CBS module reside atop of their respective CDs. The overall architecture is defined by a central helical stalk formed by antiparallel CBS modules alternatively flanked by the CDs dimers on both sides. **E** – A single *cis*-dimer 3D reconstruction focused on the flexible linker that acts as a hinge enabling substantial conformational shifts. **F** – Side view of the atomic model of a representative *cis*-dimer with bound SAO. **G** – Close-up view of the CBS module with SAO bound at the site S2 located at the interface of the rearranged RDs from both protomers. The binding pocket is primarily stabilized by a hydrophobic patch formed by helix α17 (residues 536–549) from one subunit and helices α12 (residues 431–440) and α14 (461–469) from the complementary subunit. Together with the oligomerization loop, this arrangement promotes formation of a continuous central stalk. **H** – Electrostatic surface potential map of the CBS module interface showing charge distribution contributing to RD-RD interaction of CBS module interface.

### SAM binding facilitates CBS activation via filament stacking

To further investigate the mechanism of CBS allosteric activation, we explored the structural outcomes of CBS in the presence of its native allosteric ligand SAM. Given that the SAO-bound CBS structure challenged the prior models of CBS allosteric regulation, we sought to resolve whether SAM binding induces a comparable or distinct conformational state. Initial cryo-EM micrographs revealed that SAM binding triggered a reorganization of the *trans*-basal CBS filament into a configuration reminiscent of the SAO-bound *cis*-basal CBS filament (**Figure 5A**). In this minor population, SAM was bound at the canonical S2 site within the RDs, similar to SAO (**Figure 5B**). However, the major population captured in the presence of SAM was a novel filamentous assembly resembling two *cis*-basal filaments stacked in a zipper-like fashion (**Figure 5C** and **Supplementary figure 9D**). Due to the considerable size variation and conformational flexibility of these stacked assemblies, standard symmetry-based approaches were insufficient. Instead, multiple rounds of focused local refinements in combination with iterative helical reconstructions targeting individual substructures including peripheral CDs, RDs forming the central stalks, and stacked layers of CDs were employed. This strategy enabled us to reconstruct a composite model of the full assembly at a global resolution of 7-10 Å (**Figure 5D**).

**Figure 5.**
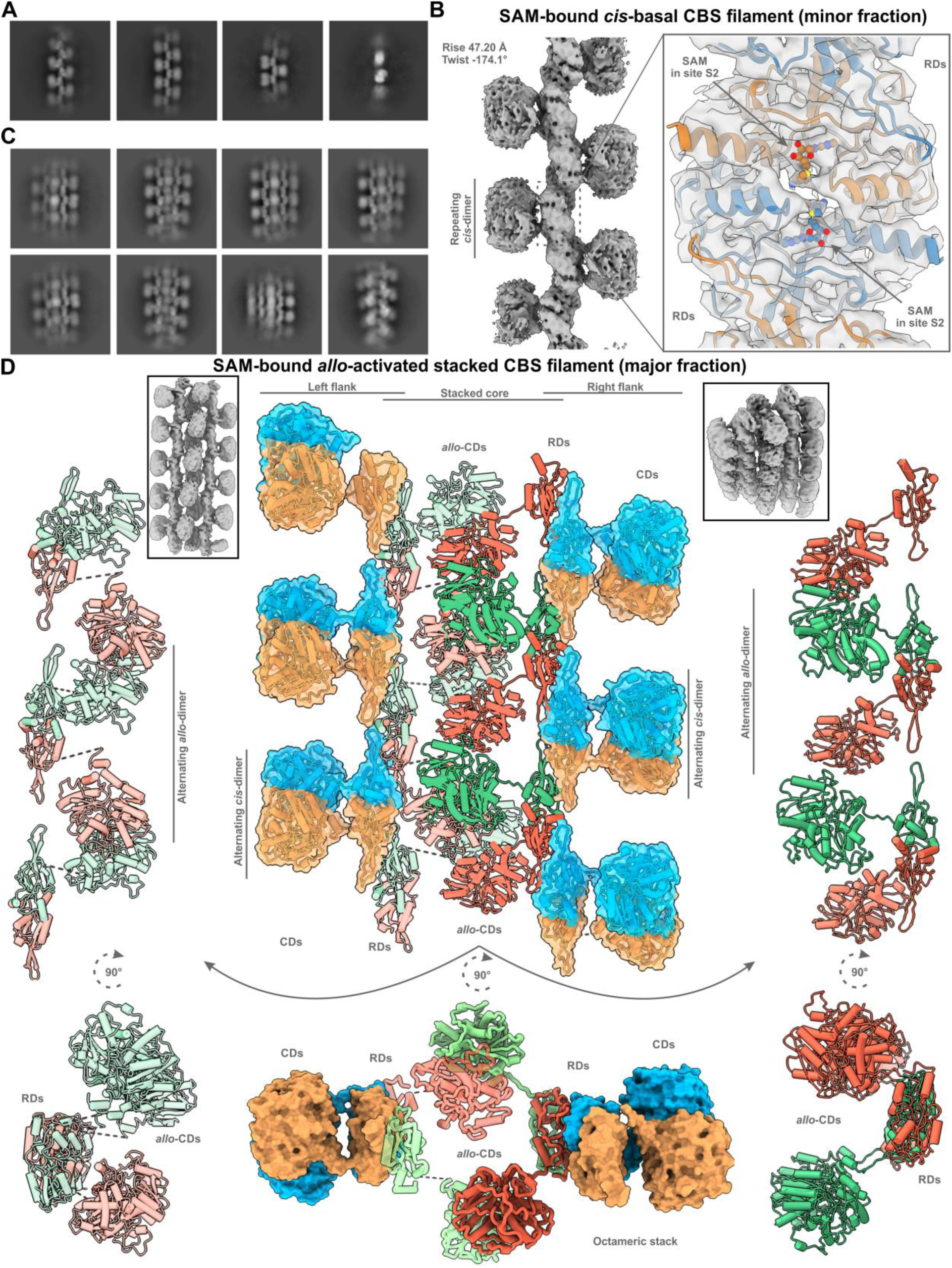
Structural characterization of SAM-induced CBS filamentous assemblies. **A** – Representative 2D class averages of SAM-induced CBS filaments representing a minor population, displaying a morphology similar to the SAO-bound *cis*-basal CBS filaments. **B** – Helical cryo-EM reconstruction of the SAM-bound *cis*-basal CBS filament essentially identical to that of SAO-bound *cis*-basal filament. Zoom-in view of the central stalk revealing SAM-bound at the S2 site within the CBS module. **C** – Representative 2D classes of the predominant population of SAM-induced CBS filaments revealing the formation of highly compact, stacked assemblies distinct from both *trans*-basal and *cis*-basal states. This unique arrangement is referred to as the *allo*-activated stacked CBS filament. **D** – Detailed architectural breakdown of the SAM-induced *allo*-activated stacked CBS filament. The inset on the left displays the overall 3D helical reconstruction of the SAM-bound *allo*-activated CBS assembly resolved at a global resolution of 7-10 Å. The inset on the right shows that this assembly exhibits a subtle right-handed helical twist, which is proposed to contribute to enhanced structural stability and dense packing. The entire quaternary structure can be broadly divided into the central stacked core and two flanking regions. The central core region comprises a unique arrangement of *allo*-dimers (CBS protomers shown in alternating dark and light red and green colors), in which the catalytic domain (CD) dimers are contributed from protomers distinct from those forming the regulatory domain (RD) CBS module, unlike the canonical *cis*-dimer arrangement. Each left- and right-side flanking region contains *cis*-dimers, where CBS modules are aligned in alternating fashion with those of the core *allo*-dimers, forming two RD stalks via oligomerization loop-mediated interactions. To better illustrate the core architecture, two separate panels (the left and the right) depict the molecular details of the arrangement. The bottom panel (top-down perspective) highlights the intricate interweaving of CDs and RDs forming the octameric building block composed of two *cis*-dimers and two *allo*-dimers. This structural arrangement underlies the allosteric activation of CBS by SAM and represents a distinct quaternary configuration not seen in basal filament states.

The major SAM-induced CBS assembly exhibited a distinctive quaternary conformation. A subtle right-handed helical twist of the stack was observed (**Figure 5D inset**) likely contributing to the overall structural stabilization. Notably, within the central region of the stacked assembly, CD dimers were alternately contributed by the two opposing filaments and were slightly displaced outward to accommodate spatial constraints. Further analysis revealed that these centrally located CD dimers were structurally distinct from the peripheral CDs, which retained the *cis*-dimer arrangement as seen in the SAO-bound or minor SAM-bound structures (**Figure 4C-D&5A-B**). Strikingly, the CBS dimer in the stacked core showed a novel association, wherein the CBS module and CD dimer were assembled from the protomers belonging to the opposing filaments. We refer to this novel configuration as the *allo*-CBS dimer (**Supplementary figure 12B**). Each stacked unit was composed of two *cis*-dimers and two *allo*-dimers, forming an octameric assembly that extended into filaments through interactions mediated by the oligomerization loop (**Figure 5D**). Beyond its structural novelty, the *allo*-dimer likely contributes directly to the CBS activation by creating a catalytically favorable interface absent in *trans*-basal and *cis*-basal CBS filaments. Despite the moderate resolution limiting detailed interpretation of the catalytic site, the unique architecture of the central CD stack provides a plausible structural basis for SAM-induced activation. The observed inter-filament association, mediated by the unique *allo*-dimers, appears to be essential for forming the catalytically competent state, which we define as the *allo*-activated stacked CBS filament.

The co-existence of multiple quaternary states raised key questions about their reversibility and physiological relevance. Enzymatic assays demonstrated that the CBS activation by SAM is reversible: a titration of SAM up to 200 µM resulted in a half-maximal activation constant (k_act_) of 7.7 µM. Conversely, pre-incubation with 200 µM SAM, followed by a gradual dilution led to CBS deactivation with a k_deact_ of 12.1 µM (**Figure 6A**). This reversibility is consistent with the dynamic nature expected for metabolically regulated allosteric enzymes. To elucidate the structural transition pathway, we employed a Ni-NTA magnetic bead immobilization assay. In the absence of SAM, His-tagged CBS remained stably immobilized on the beads. Upon addition of 500 µM SAM, a rapid release of CBS into the solution was observed, followed by a gradual decline (**Figure 6B**). These results suggest that SAM binding promotes disassembly of the *trans*-basal filament, followed by a facilitation of reassembly into *cis*-basal and *allo*-activated stacked CBS filamentous conformations. To further investigate the biological relevance of SAM-induced stacking, we monitored endogenous CBS stability in hepatoma HEP3B cells under variable SAM conditions (**Figure 6C**). Under basal conditions, CBS exhibited a half-life of 7.3 hours. This increased to 11.5 hours, a substantial 58% increase, after 1 mM methionine supplementation raising the intracellular SAM concentrations. In contrast, the inhibition of SAM synthesis using Met adenosyltransferase inhibitor AGI-43192 (10 µM) reduced the CBS half-life by 26% to 5.4 hours compared to the basal conditions. Together, these findings support a model in which SAM triggers CBS activation through stacking mediated by the unique *allo*-dimer complemented by filamentation mediated by the oligomerization loop of CBS WT. This transition not only enhances enzymatic activity but also stabilizes the protein within the cellular environment.

**Figure 6.**
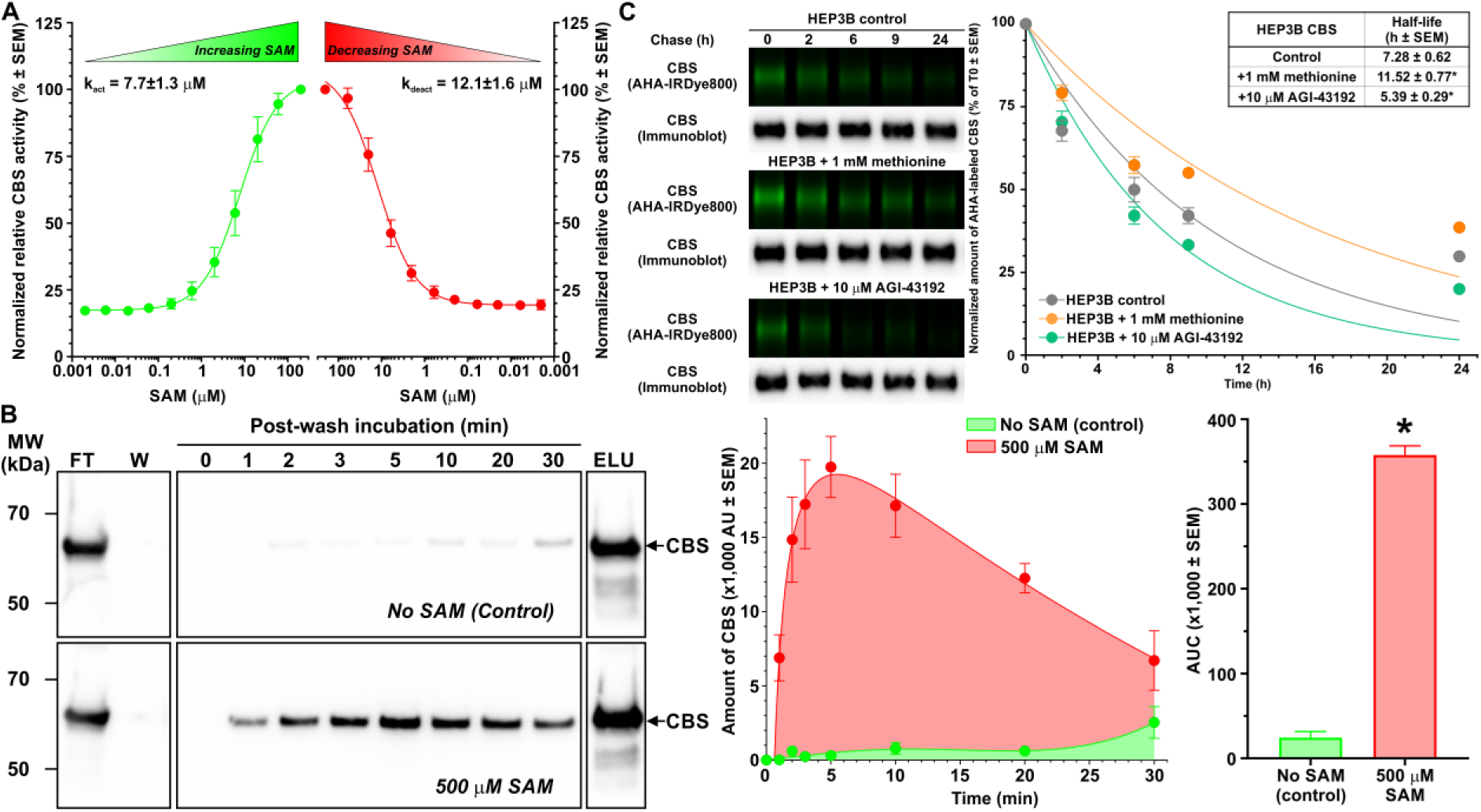
Biochemical characterization of CBS allosteric regulation by SAM. **A** – Relative CBS activities in the presence of either increasing SAM up to 200 µM (green) or decreasing SAM content (red) illustrate the reversibility of CBS activation by SAM. **B** – The SDS-PAGE Western blot (left) of major fractions (FT = flow-through, W = wash, ELU = elution) that were collected during an immobilization assay, monitoring CBS filament dynamics at different time points. In the presence of 500 µM SAM, a marked dissociation of CBS filaments was observed, followed by reassociation into the *allo*-activated stacked CBS filament. Quantification (right) illustrates this ligand-induced disassembly and reassembly process. **C** – Cellular turnover of the endogenous CBS in HEK293 cells is regulated by the intracellular content of SAM. Low SAM levels achieved by inhibition of MAT using 10 µM AGI-43192 resulted in a significantly reduced CBS half-life compared to control conditions (standard medium containing 200 µM methionine), while high levels of SAM achieved by 1 mM methionine supplementation of the culture medium substantially stabilized and increased CBS half-life. Representative gels and blots are shown on the left, were used to quantify CBS decay kinetics (right), with corresponding half-lives calculated (inset) under each condition.

## Discussion

Since its discovery and initial characterization, the oligomeric nature of human CBS has been a subject of considerable debate. Early studies variably described CBS as a heterotetramer or homodimer before converging on the widely accepted model of a homotetramer composed of four 63 kDa subunits [16–19]. However, recent findings from McCorvie and colleagues [7] and our current cryo-EM data redefine CBS architecture showing it to be a filamentous protein with repeating dimeric and tetrameric structural units (**Figure 1**). Our findings represent a transformative advancement in understanding of CBS structure-function relationships, directly linking filamentation to the enzyme’s stability, activity, and cellular turnover.

Based on our structural and biochemical analysis, we propose a dynamic model of CBS oligomerization governed by the morpheein paradigm of allosteric regulation, in line with a model previously proposed by Jaffe [20]. According to this model, homo-oligomeric proteins transition reversibly between multiple quaternary states with distinct properties. These transitions require dissociation into lower-order species and a subsequent conformational change before reassembly. This framework explains the multifaceted kinetic behaviors and regulatory features observed in CBS. Filamentous human CBS exists in at least three distinct states: the ligand-free *trans*-basal filament (**Figure 1**), the non-activating ligand-bound *cis*-basal filament (**Figure 4**) and the activating ligand-bound *allo*-activated stacked filament (**Figure 5**). Importantly, our findings show that both the *trans*-basal and *cis*-basal filaments exhibit basal catalytic activity, while only the *allo*-activated stacked filament formed through previously unseen SAM-mediated *allo*-dimer association shows catalytic activation. This rules out the previous model proposing a gradual *in situ* RD rearrangement without filament disassembly [7], which is incompatible with the structural homogeneity observed in our reconstructions. In our updated model depicted in **Figure 7**, CBS dimers associate in an allosteric ligand-dependent mode. In the absence of ligands, RDs are unable to form stabilized CBS modules and instead undergo swapping with CDs of the opposing subunit, generating the *trans*-dimer. These *trans*-dimers assemble into *trans*-basal filaments via the oligomerization loop (residues 516–525) (**Figure 1**). Binding of non-activating ligands, such as SAO or SAH, facilitates the formation of ligand-stabilized rearrangement of CBS modules into a *cis*-dimer configuration, which further oligomerizes into a straight *cis*-basal filament (**Figure 4**). In contrast, SAM induces the formation of a distinct *allo*-dimer that, along with *cis*-dimers, builds an octameric stack. This stack is extended into the catalytically competent *allo*-activated stacked filament through oligomerization loop-mediated interactions (**Figure 5**). The central *allo*-CDs in this assembly adopt a novel geometry that likely enhances catalytic activity through effects on local crowding, diffusion, and protein dynamics [21]. This hierarchical assembly requiring initial dissociation followed by reassociation aligns well with a proposed morpheein framework. Importantly, stacking and filamentation appear to occur concurrently rather than sequentially, with the octameric stack acting as a structural seed that nucleates the formation of the *allo*-activated stacked filament (**Figures 5**). This also explains why CBSΔ516-525, despite the lack of the oligomerization loop and inability to form filaments, is still activated by SAM via a transient formation of the octameric stack.

**Figure 7.**
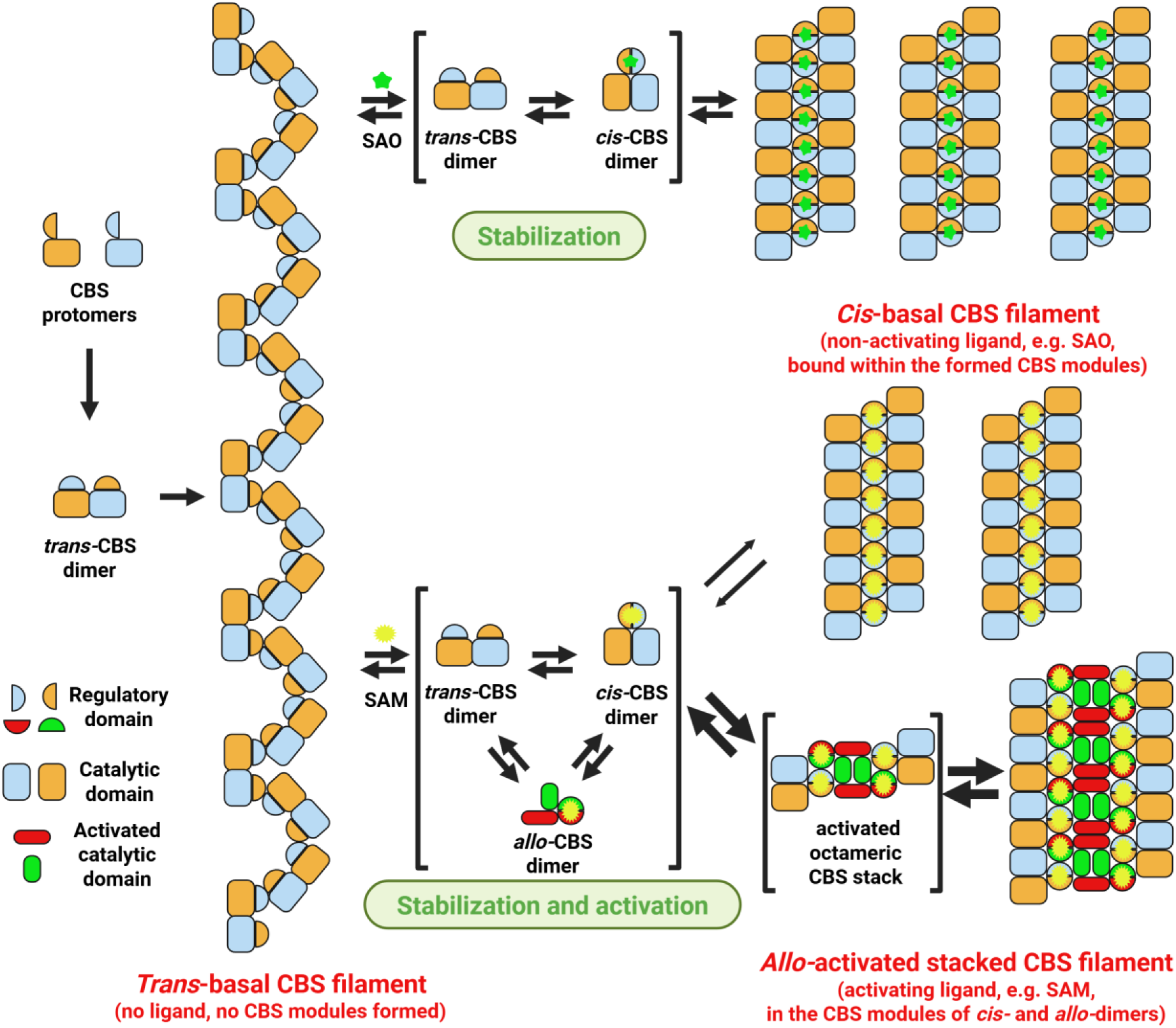
A morpheein-based model of CBS oligomerization and allosteric regulation. Nascent human CBS polypeptides (depicted in orange and blue) first assemble into *trans*-CBS dimers, where the regulatory domains (RDs, shown as half circles) are swapped and interact with the catalytic domains (CDs) of the opposing subunit. Oligomerization of these *trans*-dimers, mediated by the oligomerization loop (residues 516-525), results in the formation of the *trans*-basal CBS filament, which represents the CBS enzyme’s basal, ligand-free state. Binding of a non-activating allosteric ligand, such as adenosylornithine (SAO; green star), results in the dissociation of the *trans*-basal CBS filament, conformational rearrangement of the CBS subunits yielding *cis*-CBS dimers characterized by the formation of a ligand-stabilized CBS module between the RDs of the complementary subunits within the dimer. Subsequent oligomerization of these *cis*-dimers mediated by the same oligomerization loop gives the *cis*-basal CBS filament, characterized by increased stability and retains the basal enzymatic activity. Binding of an activating allosteric ligand, such as S-adenosylmethionine (SAM; yellow star), triggers a similar dissociation of the *trans*-basal CBS filament, but promotes a distinct conformational outcome: formation of *allo*-CBS dimers. These dimers exhibit a CBS module stabilized by RD interactions similar to those observed in *cis*-dimers; however, their CDs form dimers with CDs from adjacent *allo*-dimers, enabling a unique inter-dimer arrangement. Together with SAM-bound *cis*-dimers, these assemble into an octameric CBS stack composed of two *cis*-dimers and two *allo*-dimers. This stacked octamer serves as a nucleation seed for the formation of the *allo*-activated stacked CBS filament, a higher-order assembly characterized by high enzymatic activity and structural stability. The central CD dimers in this filament (shown in red and green) adopt a distinct conformation not seen in other CBS assemblies. The stacking and the filamentation occur concurrently, facilitated by the oligomerization loop (residues 516-525). Additionally, a minor population of SAM-bound *cis*-dimers associates independently of the *allo*-dimers yielding SAM-bound *cis*-basal CBS filament similar to those induced by the non-activating ligands like SAO. Human CBS enzyme thus exists in a dynamic equilibrium between these distinct quaternary states, modulated by both CBS concentration and the nature and availability of the allosteric ligand.

Our model reconciles the binding stoichiometry and affinities of CBS-ligand interactions reported by isothermal titration calorimetry [2, 5–7]. Filamentous CBS WT showed biphasic binding curves for SAM consistent with two types of binding sites [2], whereas engineered dimeric CBSΔ516-525 only exhibited a single site binding [6]. We interpret these differences as a reflection of structural transitions, i.e. dimer rearrangement upon binding and formation of higher-order oligomeric assemblies, that depend on both the character of the allosteric ligand and the presence of the oligomerization loop. Additional supporting evidence comes from studies employing differential scanning calorimetry, which showed that SAM binding substantially stabilized the RD, an effect significantly less prominent with non-activating ligands, such as SAH and SAO [2, 5, 6].

Although earlier interpretations designated the *cis*-dimer as the activated form based on comparisons to structures of *Drosophila melanogaster* CBS and human CBSΔ516-525 E201S variant [3, 22], our findings reveal that activation of human CBS is driven only when SAM mediates formation of the *allo*- and *cis*-dimers, which yield activated octameric stack, while the intact oligomerization loop facilitates formation of the *allo*-activated stacked CBS filament. These findings suggest that SAM not only stabilizes the CBS module, but – unlike non-activating ligands, such as SAH and SAO – also drives a specific spatial reconfiguration required for activation.

Filamentation of CBS may not be conserved across all species. Our recent work on mouse CBS, which lacks filamentation capability due to a longer oligomerization loop (similar to CBS ISO2 variant), suggests this feature may be species-specific [23]. The biological roles of filamentation in CBS likely mirror those seen in other metabolic enzymes [11], such as providing regulatory flexibility [24, 25], enabling compartmentalization [26] or protecting against degradation [27]. Indeed, our data show that non-filamentous dimeric CBS constructs exhibit significantly shorter cellular half-lives and altered sub-cellular localization compared to the filament-forming CBS WT (**Figures 2** and **6**).

The proposed morpheein-based model of CBS regulation has significant pathophysiological implications. Homocystinuria (HCU) is an inborn error of metabolism and conformational disorder resulting from protein misfolding, rapid degradation and/or impaired regulation of pathogenic CBS variants [12, 28–31]. Mutations that interfere with rearrangement of CBS dimers, their assembly into higher order oligomers, impair filamentation or promote aggregation may drive the disease phenotype. Therapeutic strategies targeting CBS filamentation, either by stabilizing functional assemblies or promoting proper allosteric transitions, could offer new therapeutic avenues. Particularly, SAM analogs have been hypothesized to act as CBS-specific pharmacological chaperones rescuing enzyme’s oligomerization, stability and activity [5, 31, 32]. Anticancer drug taxol, which allosterically inhibits the assembly and disassembly of microtubules by binding tubulin [33], or morphlock-1, which inhibits transition of porphobilinogen synthase from inactive hexamers to active octamers [34], set a precedent for feasibility and successful development of novel pharmacological therapies regulating CBS. These may extend beyond HCU into conditions with dysregulated CBS including wound healing, angiogenesis, Down syndrome and various types of cancer [35–37].

In conclusion, our findings redefine CBS as a filamentous morpheein and establish assembly, stacking and conformational plasticity as central features of its regulatory landscape. This paradigm shift opens exciting new possibilities for understanding and therapeutically modulating CBS function.

## Materials and Methods

### Preparation of human CBS protein

Full-length human CBS was expressed and purified as described previously [15, 38] with a few modifications. Briefly, *E. coli* Rosetta2 (DE3) cells transformed by pGEX-6P1-hCBS construct were cultured using twelve 2.8l Fernbach flasks each containing 1l of LB medium supplemented with 0.001% thiamine.HCl, 0.0025% pyridoxine.HCl, 0.15 mM FeCl_3_, 3% ethanol, 0.3 mM delta-aminolevulinic acid and 100 µg/ml ampicillin. The cells were grown at 30°C shaking at 275 rpm until the OD_600_ reached ∼0.75. The expression of CBS was induced by the addition of 0.5 mM IPTG. After overnight expression, cells were harvested by centrifugation at 8,000 rpm, 4°C for 5 min, pooled, washed with ice-cold 1xPBS and immediately frozen at -80°C. Later, the cells were resuspended in cold lysis buffer (50 mM sodium phosphate pH 7.4, 300 mM NaCl, 1 mM DTT, 1% Triton X-100, 0.1 mM PLP, 1x Sigma Protease inhibitor cocktail) using Dounce homogenizer and lysed with 2 mg/ml lysozyme while rocking at 4°C for 1 hour. After sonication to decrease viscosity of the suspension, the insoluble fraction was separated by centrifugation at 21,000 rpm, 4°C for 30 min. The GST-CBS fusion protein was purified by loading the clarified lysate on the GST Sepharose column, washed with 10 mM sodium phosphate pH7.4, 500 mM NaCl, 1 mM DTT and eluted with a wash buffer containing 20 mM reduced glutathione. The GST fusion partner was cleaved off by 0.5 U HRV3C protease per milligram of protein at 4°C overnight followed by a separation of GST and CBS using 15-300 mM potassium phosphate pH 7.2 gradient on DEAE Sepharose column. The CBS-rich fractions were pooled, concentrated and buffer exchanged into 20 mM HEPES pH 7.4, 1 mM TCEP and 0.01% Tween 20 using Amicon YM-100 membrane. Purified CBS at concentrations typically ranging 10-20 mg/ml was stored at -80°C in small (20-50 µl) aliquots.

### Cryo-EM

Four separate cryo-EM datasets were prepared and analyzed: (1) CBS WT in its basal state (*trans*-basal CBS), (2) CBS bound to its substrate L-serine (Ser-*trans*-basal CBS), (3) CBS bound to allosteric ligand sinefungin (adenosylornithine, SAO) (SAO-*cis*-basal CBS) and (4) CBS bound to SAM (SAM-*allo*-activated stacked CBS).

For the *trans*-basal CBS dataset, concentrated CBS WT was diluted to 2.2 mg/ml in a 20 mM HEPES pH 7.2, 100 mM NaCl and 1 mM β-mercaptoethanol. The sample was applied to freshly plasma-cleaned gold Quantifoil R 1.2/1.3 grids (300 mesh) and vitrified using a Vitrobot Mark IV (ThermoFisherScientific) with 100% humidity at 10°C, using a blot force of 0 and blot time of 3.5-4.5 seconds. For the Ser-*trans*-basal CBS, 0.5 mM L-serine was added to CBS and incubated for 20 min on ice before grid preparation. Similarly, SAO-*cis*-basal CBS grids were prepared by incubating CBS with 1 mM SAO for 20 min. To capture SAM-*allo*-activated stacked CBS, CBS was incubated with 0.5 mM SAM for 10 min, generating samples with different stages of allosteric activation. All grids were prepared under identical controlled conditions, ensuring consistency in sample vitrification. Cryo-EM data for all four datasets were collected on Titan Krios G4 microscopes (ThermoFisherScientific) operated at 300 keV, equipped with a Falcon 4 direct electron detector operating in counting mode. For *trans*-basal CBS and Ser-*trans*-basal CBS datasets, a SelectrisX energy filter with a 10-eV slit width was used. Data acquisition was automated using EPU software (v3.5.1), utilizing aberration-free image shift (AFIS) for efficient high-throughput collection.

#### The *trans*-basal CBS (dataset 1)

For the *trans*-basal CBS conformation, a total of 4,865 movies were recorded in Electron Event Representation (EER) mode, using a calibrated pixel size of 0.72 Å and a total electron dose of 64 e⁻/Å². After careful assessment and selection based on CTF estimation and overall image quality, 3,530 movies were chosen for further processing. Initial motion correction and dose weighting were performed using the built-in tools in cryoSPARC (v4.4.1) [39]. The contrast transfer function (CTF) parameters were estimated on a per-micrograph basis [40]. For filament-based helical reconstruction, filaments were manually traced using cryoSPARC’s filament tracer module with an inter-box separation distance set at 32 Å. An initial set of approximately 1.2 million filament segments was extracted. Multiple iterative rounds of 2D classification were performed to remove false positives and damaged particles, leading to a high-quality subset containing 528,877 particles. This refined set was used for an *ab-initio* reconstruction into four classes. Two well-defined classes were selected and combined for further processing (**Supplementary figure 1**). Initial helical parameters (twist and rise) were estimated using a combination of tools, including cryoSPARC’s symmetry search utility, HELIXPLORER (https://rico.ibs.fr/helixplorer/), *ab-initio* volume reconstructions examined in UCSF Chimera [41, 42], and analysis of power spectra from elongated filament segments (**Supplementary figure 2**).

Subsequently, heterogeneous refinement with two classes was performed and the best class was selected for high-resolution helical refinement. A final set of 335,525 particles was used, yielding an initial reconstruction at 3.4 Å resolution. Further iterative refinements were carried out using C1 and D1 symmetries separately, resulting in final maps at 3.2 Å and 3.0 Å, respectively. The final refined helical parameters were a twist of –116.11° with a rise of 49.9 Å for C1 symmetry, and –116.06° with a rise of 49.97 Å for D1 symmetry, respectively. For single-particle-like (SPA-like, non-helical) processing, an independent particle-picking strategy was employed. An initial set of approximately 1.4 million particles was extracted. Extensive rounds of 2D classification were performed to clean the dataset, ultimately retaining 437,000 high-quality particles. These particles underwent an *ab-initio* 3D classification into three classes, from which two well-resolved classes were selected and merged. The merged particle set, consisting of 338,925 particles, was subjected to homogeneous refinement, non-uniform (NU) refinement, and local refinement steps [43]. Application of D1 symmetry during NU refinement and subsequent focused local refinement on the dimer–dimer interface significantly improved map quality. The resolution progressively improved from 2.89 Å (without symmetry, C1) to 2.63 Å when D1 symmetry was imposed, following final global CTF aberration correction and additional non-uniform and local refinements (**Supplementary figure 1**). The final high-resolution map allowed for detailed visualization of side-chain densities and confident atomic model building.

#### The Ser-*trans*-basal CBS (dataset 2)

For the Ser-*trans*-basal CBS (dataset 2), a total of 14,954 movie stacks were collected in EER format, out of which 13,200 movies were selected after careful evaluation of CTF estimations and overall image quality. Data collection was performed at a nominal pixel size of 0.452 Å and with a total electron dose of approximately 52 e⁻/Å² per movie, allowing for fine sampling of high-resolution features. Using the filament tracer tool implemented in cryoSPARC, initial particle picking was carried out along the filaments with an inter-particle separation of 50 Å. This approach resulted in an initial set of approximately 2.18 million filament segments. Several iterative rounds of 2D classification were subsequently performed to remove poor-quality or contaminant particles, refining the dataset to around 1.2 million particles suitable for downstream processing. Following 2D cleaning, an *ab-initio* reconstruction was performed with three initial classes. Two well-defined classes exhibiting clear secondary structure features were selected and merged for further helical reconstruction (**Supplementary figure 3**). Initial helical parameters, including twist and rise, were determined as described previously.

Approximately 1 million particles were subjected to helical refinement using both C1 and D1 symmetry constraints, yielding final resolutions of 2.68 Å and 2.70 Å, respectively. The optimized helical twist was refined to –114.6°, and the rise converged at approximately 49.9 Å. To further dissect the heterogeneous intermediates trapped during catalysis, several rounds of focused 3D classification were performed. Classification jobs with K=5, 8, and 10 classes were run after anisotropic correction to separate distinct catalytic substates. After detailed analysis, three classes comprising a combined total of 577,766 particles were merged from K=8 classification for additional focused refinement. These merged particles were also subjected to an SPA-like processing, where duplicates closer than 60 Å were excluded, followed by NU refinement and further helical refinement. This workflow yielded maps with resolutions of 2.34 Å and 2.65 Å for the major intermediate states (**Supplementary figure 3**). To further resolve local conformational differences, a final round of focused 3D classification was performed using custom masks spanning the full catalytic domain and flanking regulatory interfaces, using K=5 classes and C1 symmetry (D1 symmetry was not imposed in any of the 3D classifications). The three best 3D classes were individually refined, revealing subtle conformational rocking motions of the RDs relative to the CDs. Final SPA-like refinements on these distinct classes led to the identification of three well-resolved substrates corresponding to (i) internal aldimine CBS-Lys-PLP refined to 2.72 Å resolution, (ii) external aldimine CBS PLP-Ser refined to 2.20 Å resolution and (iii) aminoacrylate CBS-PLP-AA refined to 2.00 Å resolution (**Supplementary figure 4**). Corresponding helical refinements with D1 symmetry constraints yielded consistent maps at 2.97 Å (twist –115.7°, rise 50.4 Å), 2.77 Å (twist – 115.0°, rise 50.8 Å) and 2.54 Å (twist –115.3°, rise 49.9 Å), respectively (**Supplementary figure 5**). These final high-resolution maps allowed for unambiguous visualization of the ligand densities and precise side-chain positioning, facilitating confident atomic model building and enabling detailed mechanistic interpretation of each intermediate state.

#### The SAO-*cis*-basal CBS (dataset 3)

For the SAO-*cis*-basal CBS (dataset 3), a total of 14,300 movies were collected in EER format. Data collection was performed at the nominal pixel size was 0.52 Å, and the total electron dose was set at 50.0 e⁻/Å². Following initial motion correction and CTF estimation, 12,400 high-quality movies were retained for further processing. Particles were automatically picked using filament tracer in cryoSPARC with an inter-particle separation of ∼50 Å. After the initial extraction, approximately 2.2 million filament particles were obtained. These underwent multiple rounds of 2D classification to discard broken or low-quality segments, resulting in a refined subset of 1.1 million particles. This refined subset was subjected to two rounds of *ab-initio* reconstruction with three classes each, after which two well-defined classes were selected and merged. Further 2D classification was performed to isolate the best filament views, yielding a final particle set of 411,125 particles for helical refinement. Initial helical parameters, including twist and rise, were determined using cryoSPARC, HELIXPLORER, and real-space measurements in UCSF Chimera. Helical refinement using both C1 and D1 symmetries was then carried out, producing final maps with a helical twist of 172.4° and a rise of 48.47 Å, reaching an overall resolution of 4.2 Å (**Supplementary figures 6&7**).

To improve local detail, focused refinement on the central regulatory domain stalk was performed using a soft mask, increasing the resolution locally to 4.0 Å. Additionally, separate helical refinement focusing on two catalytic domains and adjacent regulatory domains was conducted to highlight the linker motif connecting CDs and RDs, achieving a resolution of approximately 4.8 Å (**Supplementary figures 6&7**). In parallel, an SPA-like approach was employed. Duplicate particles were removed from the helical dataset (filtered by a minimum separation threshold), followed by two further rounds of 2D classification and one *ab-initio* reconstruction. This process yielded 106,308 particles, which were subjected to NU and local refinements to better resolve one catalytic domain and two regulatory domains, clarifying inter-domain interfaces (**Supplementary figure 8**). Finally, an overall filament-focused local refinement without explicit helical symmetry (C1 symmetry imposed) was performed to validate structural consistency. No significant differences were observed when comparing maps generated using C1 and D1 symmetries, confirming the stability and homogeneity of the SAO-*cis*-basal CBS conformation.

#### The SAM-*allo*-activated stacked CBS (dataset 4)

For the SAM-*allo*-activated stacked CBS (dataset 4), two batches of movies were collected: one consisting of 3,910 movies used primarily to optimize grid preparation and enhance the number of non-aggregated assemblies, and a second larger batch of 8,670 movies. After CTF correction and initial screening, 11,510 high-quality movies were selected from a total of 12,580 movies recorded in EER format. Movies were collected at a total electron dose of 42 e⁻/Å² and a pixel size of 0.83 Å.

Filament particles were initially picked using cryoSPARC’s filament tracer tool, testing separation distances of 30 Å, 40 Å, 60 Å, and 100 Å, ultimately choosing 100 Å separation for the best capture of the stacked filament architecture. An initial set of ∼620,000 particles was extracted and subjected to multiple rounds of 2D classification to remove low-quality picks and aggregates, resulting in a refined set of 120,000 particles. *Ab-initio* reconstructions with K=4 classes were performed, and the two best classes were merged for initial helical reconstructions. Helical parameters, including twist and rise, were determined using cryoSPARC tools, real-space marker measurements in UCSF Chimera, and analysis of power spectra of elongated filaments (**Supplementary figure 9**).

Several iterative rounds of helical refinement and reconstruction were conducted, progressively restricting the alignment resolution to 12, 10, 8, and 6 Å to enhance overall map quality and minimize misalignment artifacts. C1, D1, and C2 helical symmetries were systematically tested to improve map fidelity and to verify that no artifacts were introduced by symmetry imposition (**Supplementary figure 10&11**). Subsequent two rounds of 3D classification (K=3), followed by additional 2D classification to exclude suboptimal segments, resulted in a final refined subset of 33,700 particles for helical reconstruction. The RDs stalks and outer CDs displayed an overall right-handed twist with optimized parameters of 175.6° and a rise of 48 Å, while the central stacked CDs region showed a distinct structural arrangement.

In parallel, an SPA-like analysis was also performed. From the initial extraction, ∼1.25 million particles were subjected to multiple rounds of 2D classification. A final subset of 41,174 particles, corresponding to stacked filament segments, was selected after duplicate removal using an 80 Å separation cutoff. These particles were refined through helical refinement, NU refinement, and local refinement to validate map consistency and to resolve orientations of the outer CDs observed in 2D classes. Final reconstructions using this approach achieved resolutions between 7 and 10 Å, providing improved visualization of the outer CDs and RDs stalks (**Supplementary figure 12**).

Focused classification and refinement with masks centered on the core of the filament were performed in both helical and SPA-like frameworks. Further NU and local refinements were used to attempt to resolve the diffuse central CD density and to trace the linker motifs connecting the RDs to the CDs in the center of the stack. Despite extensive efforts, the resolution of the central CD region did not improve substantially; however, these refinements clarified which RDs contribute to the central CD assembly. From the ∼8 Å maps, we concluded that the motif organization and CD dimer assembly in the central core of the SAM-*allo*-activated stacked CBS filament are distinct from those in the *trans*-basal, *cis*-basal, and outer stacked CDs.

In addition, similarly to the SAO-*cis*-basal CBS dataset, we reconstructed a SAM-bound *cis*-basal CBS conformer. Using helical reconstruction, this structure reached an overall resolution of 4 Å, which was further improved to 3.8 Å through NU, local, and SPA-like refinements (**Supplementary figure 13**). We also observed various fragmented assemblies, including isolated RD filaments, partial CDs not arranged into filaments, and incomplete tetrameric structures, highlighting the structural heterogeneity induced by SAM binding (**Supplementary figure 14**).

### Model building, refinement, and validation

For the *trans*-basal CBS and Ser-*trans*-basal CBS (datasets 1 and 2) structures, initial models were prepared by docking a crystal structure of CBSΔ516–525 (PDB# 4COO) into the cryo-EM maps using rigid-body fitting in UCSF Chimera/ChimeraX [41, 42]. The missing loop region (residues 513–527), critical for oligomerization, was manually rebuilt in Coot based on visible cryo-EM density and known secondary structure predictions [44].

Multiple copies of the refined CBS protomer were then placed into the helical assemblies guided by the determined symmetry parameters. Initial fits were followed by iterative rounds of real-space refinement using Phenix to optimize geometry, improve map-to-model correlation, and minimize steric clashes [45, 46]. Flexible fitting steps were performed using Coot allowing local conformational adjustments to better accommodate subtle variations within the filament interfaces [44].

For the SAO-*cis*-basal CBS (dataset 3), models of the RD and CD were fitted individually into focused maps. The CBS module containing the RD in *cis* conformation was modeled based on the SAM-bound CBSΔ516–525 E201S structure (PDB# 4UUU) as a template. SAM analog binding sites were initially modeled using known ligand densities, and side chain positions were refined to match the local cryo-EM density.

For the SAM-*allo*-activated stacked CBS (dataset 4), modeling involved fitting separate RDs into the central stalks and CDs into the peripheral densities. The RD models from the basal structures were used as templates and flexibly adjusted in ChimeraX and Coot [41, 42, 44]. Due to partial flexibility and lower local resolution of the CDs, models of the CD (PDB# 4PCU) were manually docked into focused refined maps and adjusted iteratively. The novel inter-filament interfaces forming the *allo*-dimer arrangement were carefully built and validated using visual inspections of cross-correlation in UCSF ChimeraX [42].

Ligands (SAM, PLP, and serine) were manually docked in Coot into corresponding densities and refined in Phenix real-space refinement [45, 46]. Iterative validation of fit was performed using map-model FSC curves, local resolution-based adjustments, and MolProbity validation tools to monitor geometry (Ramachandran plot, rotamer outliers, clashscore) [47].

Final models were subjected to comprehensive validation with MolProbity to assess overall stereochemistry, hydrogen bonding networks, and sidechain rotameric states [47]. The refined models displayed acceptable overall geometry with minimal outliers and were consistent with the cryo-EM densities across the entire filament structures.

### Preparation of CBS-expressing cell lines

The HEK293A cells lacking CBS (CBS knockout) were generated by the CRISPR-Cas9 approach as detailed elsewhere [12]. Stable cell lines expressing various human CBS constructs (ISO1, ISO2, Δ516-525 and CBS45) were prepared as described previously [12]. Briefly, codon-optimized human CBS WT (ISO1, Uniprot# P35520-1) and isoform 2 (ISO2, Uniprot# P35520-2) carrying myc-tag and 6xHis-tag at their N- and C-termini, respectively, were synthesized by Genscript and subcloned using pLentiCMVBlast-empty (w263-1) plasmid (cat# 17486, Addgene) directionally into unique SalI and XbaI restriction endonuclease sites. The CBSΔ516-525 and CBS45 (CBSΔ414-551) were prepared by site-directed mutagenesis by Genscript using CBS WT (ISO1) plasmid as a template. All constructs were verified by DNA sequencing (Genscript). Supplied plasmids were used for the generation of lentiviral particles, which were used to transduce HEK293A CBS KO in the presence of 6 μg/ml protamine sulfate. Blasticidin S (45 μg/ml) was added to the culture 72 h after transduction to select for cells carrying CBS construct.

### Determination of CBS half-life

Cellular turnover of overexpressed CBS constructs was determined by using non-radioactive method based on the incorporation of L-azidohomoalanine (AHA), a biorthogonal analog of Met, as described previously [12] with a few modifications. The cells were grown to ∼80% confluency in a six-well plate. The cells were then washed with Met-deficient DMEM containing 10% dialyzed FBS, 2 mM GlutaMax, 1 mM pyruvate and 200 μM cystine hydrochloride followed by live-labeling for 2 h (pulse) in a Met-deficient DMEM supplemented with 75 μM AHA. The AHA pulse was terminated by removing the labeling medium and washing with a complete standard medium, then incubated in the same medium for up to 24 h at 37°C. Immediately after the pulse (0 h) and at designated timepoints during the chase period (2, 4, 6, 9, 12 and 24 h), the cells were harvested, lysed and the lysates analyzed as described previously [12].

The half-life of endogenous CBS under different metabolic conditions was determined similarly as described above with the following modifications. The HEP3B cells were seeded at 30% confluency and cultured in 100 mm dishes for 48 h. After the 4 h labeling pulse with 100 μM AHA, the AHA-containing Met-deficient DMEM medium was replaced with the complete low-glucose DMEM medium (control) or the same medium supplemented with 1 mM Met or 10 µM AGI-43192. Cells were cultured for up to 24 h after the AHA pulse and harvested at designated timepoints (0, 2, 6, 9 and 24 h).

### Confocal imaging

The cells were seeded into an 8-well cell culture chamber glass slide with a removable frame at a density of 80,000 cells/well. After overnight incubation, the cells were washed, fixed with 4% PFA for 15 minutes at room temperature (RT), washed three times, then permeabilized in PBS containing 0.2% Triton X-100 for 10 min at RT and washed three times before being incubated in the 10% goat serum for 10 min at RT. Cells were then washed and incubated with anti-CBS monoclonal antibody (D8F2P) at a concentration of 1:500 overnight at 4°C. The following day, cells were washed three times with PBS and incubated with the Alexa Fluor Plus 568-conjugated anti-rabbit secondary antibody at a 1:1,000 dilution for 1 hour at RT. Nuclear stain DAPI (5 µg/ml) was added for the last 5 minutes of the incubation. Cells were then washed three times with PBS and the cover slip was mounted using Prolong Gold antifade reagent. Slides were dried overnight in the dark and stored at -20°C until further use. Leica TCS SP5 confocal microscope equipped with a motorized conventional Galvo stage was used for the visualization of cells in a Z-stack. Several optical sections were acquired along the Z-axis at 1 μm step size using 40x magnification. Series of images were captured using an image format of 1024×1024 pixels and 200 Hz scan speed. DAPI was excited using a 405 nm UV laser, while CBS was visualized using a 561 nm DPSS laser. Fluorescence emission was recorded at 419–474 nm (DAPI) and 584-683 nm (Alexa Fluor 568) in a sequential mode. In each chamber, images from two randomly selected regions were acquired and all experiments were independently performed five times. This resulted in 10 regions of interest for each group with more than 100 cells per group. Image reconstruction was completed from series of Z-stack images using Imaris v10.0.1 software. Identical parameters for fluorescence intensity settings were applied for each group in all the four investigated groups. Images are shown as a single layer from the middle part of the stack and as a 3D reconstructed z-stack in series of all images. The corrected total CBS fluorescence was quantified by Fiji/ImageJ software using the CTCF formula: CTCF = Integrated density - (Area of selected cell x Mean fluorescence of a background). Results are shown as mean ± standard errors of the mean (SEM), from five independent experiments. Graphpad Prism 8 software was used for statistical analysis using one-way ANOVA followed by the Tukey’s post-hoc test. The value of *p*<0.05 indicated statistical significance denoted by asterisk (*).

### Protein electrophoresis and Western blot analysis

Cell lysates were mixed with either 2x Novex Tris-glycine Native sample buffer for native electrophoresis or 4x Bolt LDS sample buffer containing sample reducing agent for reducing denaturing electrophoresis. Denaturation of protein samples was achieved by heating at 95°C for 10 min. Native proteins were resolved in NativePAGE 4–16% Bis–Tris gels in 1x NativePAGE running buffer at 4°C, while reduced, denatured proteins were separated in NuPAGE 4–12% Bis–Tris gels in 1xNuPAGE MES SDS running buffer. Subsequently, proteins were transferred onto the PVDF membrane using an iBlot 2 gel transfer device (Invitrogen). Membranes were blocked in 5% non-fat milk in TBST (Tris buffered saline supplemented with 0.1% Tween 20) for 1 h at RT, followed by incubation with the primary antibody while gently agitating either for 1 h at RT or overnight at 4°C. Primary antibodies were diluted in TBST containing 5% BSA anti-CBS (1:2,000, CST# 14782) or anti-beta-actin (1:5,000; Sigma# A1978). After washing three times with TBST for 5 min, blots were incubated with anti-rabbit or anti-mouse IgG HRP-conjugated secondary antibody (1:5,000; CST# 7074 and 7076, respectively) in TBST containing 5% non-fat milk for 1 h at RT. After washing with TBST for 5 min twice and TBS for 5 min twice, the proteins were visualized with Radiance Plus chemiluminescence substrate using the Azure Imaging System 300. Captured images were analyzed using the Fiji/ImageJ and plotted by Graphpad Prism.

### CBS activity assay

CBS activities in native gels and cell lysates were determined as described previously [12]. Briefly, cell lysates (10 µg/lane) were resolved on native PAGE and the gel was developed in a staining solution (100 mM Tris.HCl pH 8.0, 20 mM cysteine, 50 mM beta-mercaptoethanol, 100 μM PLP, 200 μM lead acetate) in the absence or presence of 200 μM SAM. The gels were gently shaking at 37°C until the active CBS bands became apparent. The reaction was terminated after the same amount of time for both conditions by placing the gels into 7% acetic acid. The gels were scanned, and the CBS activity was quantified by densitometry using ImageJ.

CBS activity in the lysates or purified proteins was determined using H_2_S-specific fluorescent probe 7-azido-4-methylcoumarin (AzMC) as described previously [48]. Briefly, the reaction (200 μl total volume) containing 50 mM Tris.HCl pH 8.6, 5 μM PLP, 10 μM AzMC,10 μg of cell lysate or 1 µg of purified CBS was initiated after 10 min equilibration at 37°C with a mixture of substrates yielding final concentrations of 2 mM cysteine and 500 μM homocysteine. The assays containing variable final concentrations of SAM were followed for 90 min at 37°C recording fluorescent intensities in 5 min intervals (excitation 365 nm, emission 450 nm) Spectramax M5 microplate reader. All assays were done in three replicates and the data were analyzed using Microsoft Excel and Graphpad Prism software.

To investigate reversibility of CBS allosteric activation by SAM, the full-length human CBS (10 mg/ml) incubated at room temperature for 10 min in 100 mM Tris.HCl pH 8.6 in the absence and presence of 200 µM SAM. Subsequently, standard CBS activity assay using AzMC was performed as described above in the increasing or decreasing final concentration of SAM equal to 200, 60, 20, 6, 2, 0.6, 0.2, 0.06, 0.02, 0.006, 0.002 and 0 µM. The mixtures were incubated at 37°C for 10 minutes prior addition of substrates (0.5 mM Hcy and 2 mM Cys final), which started the reaction.

### Dissociation of CBS filaments

Fifty µL slurry of Ni^2+^-chelated magnetic beads (Genscript) was washed 3x with PBS containing 0.1% Tween20. Subsequently, 300 µl of 500 µg/mL full-length human CBS containing permanent C-terminal 6xHis tag was mixed with the washed beads and allowed to bind while gently mixed at room temperature for 1 hour. Afterward, the unbound portion was removed by washing the beads 3x with PBS containing 0.1% Tween20 followed by an additional wash by PBS. The suspension was then divided into two tubes and the beads were collected at the tube wall using a magnet. The beads were resuspended in 300 µl PBS, pulled by the magnet and 30 µl aliquot of the supernatant was removed to serve as the initial time zero sample. Then, 30 µl of PBS or 5 mM SAM (to achieve 0 or 500 µM SAM final concentration) were mixed with the beads and the supernatants were collected after 1, 2, 3, 5, 10, 20 and 30 minutes ensuring that the beads are pulled by the magnet to the wall of the tube and not disturbed. After the final timepoint, the beads were eluted in 1x LDS Sample buffer. All collected samples were subsequently analyzed using SDS-PAGE and Western Blot (WB) to assess kinetics of CBS filament dissociation.

## Data availability

All cryo-EM density maps and corresponding atomic coordinates generated in this study have been deposited in the Electron Microscopy Data Bank (EMDB) and Protein Data Bank (PDB). The *trans*-basal CBS filaments in the absence of substrates or allosteric ligands are available under accession codes EMD-XXXXX/PDB YYYY, EMD-XXXXX/PDB YYYY, and EMD-XXXXX /PDB YYYY. Serine-bound *trans*-basal CBS is deposited as EMD-XXXXX/PDB YYYY, EMD-XXXXX/PDB YYYY, EMD-XXXXX/PDB YYYY, and EMD-XXXXX/PDB YYYY. The SAO-bound *cis*-basal CBS filaments are deposited as EMD-XXXXX/PDB YYYY, EMD-XXXXX/PDB YYYY, EMD-XXXXX/PDB YYYY, and EMD-XXXXX. The SAM-bound *allo*-activated stacked CBS filaments are deposited as EMD-XXXXX/PDB YYYY and EMD-XXXXX/PDB YYYY. In addition, SAM-bound *cis*-basal CBS filament is deposited as EMD-XXXXX/PDB YYYY. A complete list of all datasets and accession codes is provided in **Table 1**.

## Acknowledgments

TM acknowledges support from the University of Fribourg Research pool (22-15) and the Swiss National Science Foundation (10.001.133). This work was supported by the Swiss National Science Foundation (SNF Grants CRSII5_177195). We thank Alexander Myasnikov, Bertrand Beckert, Sergey Nazarov and Emiko Uchikawa from Dubochet Center for Imaging (DCI), Lausanne, for their cryo-EM support.

## Conflict of interest statement

None

## Authors’ contribution

IM performed all cryo-EM sample preparation, data acquisition, structure determinations, model building, refinement and mechanistic interpretations. EM performed most cellular, biochemical and imaging studies and analyzed data. TMP performed selected biochemical studies and analyzed data. LJ assisted in confocal imaging studies. KA generated stable HEK293 cell lines. FJA generated preliminary results in negative stain. CS and HS provided conceptual input, instrumentation and financial support. TM conceptualized and directed the entire project and prepared CBS proteins. TM and IM prepared the final figures and wrote the original draft of the manuscript. All authors reviewed, commented and/or edited various versions of the manuscript and approved its final form.

## Notes

### Competing Interest Statement

The authors have declared no competing interest.

