## Supplemental figures 1-14 for "Structural basis for a filamentous morpheein model of human cystathionine beta-synthase"

**Supplementary figure 1. Cryo-EM image processing workflow for structural elucidation of human CBS in the absence of substrate and allosteric ligands.** **A** – Representative electron micrograph showing long filamentous assemblies of WT-CBS. **B&C** – Representative 2D class averages (**B**) and 3D classification density maps (**C**) obtained using the helical reconstruction approach. **D&E** – Representative 2D class averages (**D**) and 3D classification maps (**E**) obtained using the single-particle analysis (SPA) approach. **F** – Final D1-symmetrized reconstruction of the *trans*-basal CBS filament from the SPA shown with local resolution map and the corresponding Fourier shell correlation (FSC) curve, as well as particle orientation distribution (**I**). **G** – Final helically reconstructed map of the *trans*-basal CBS filament at 3.0 Å resolution, shown with local resolution estimation and the corresponding FSC curve (**H**).

### The *trans*-basal CBS filament (dataset 1)

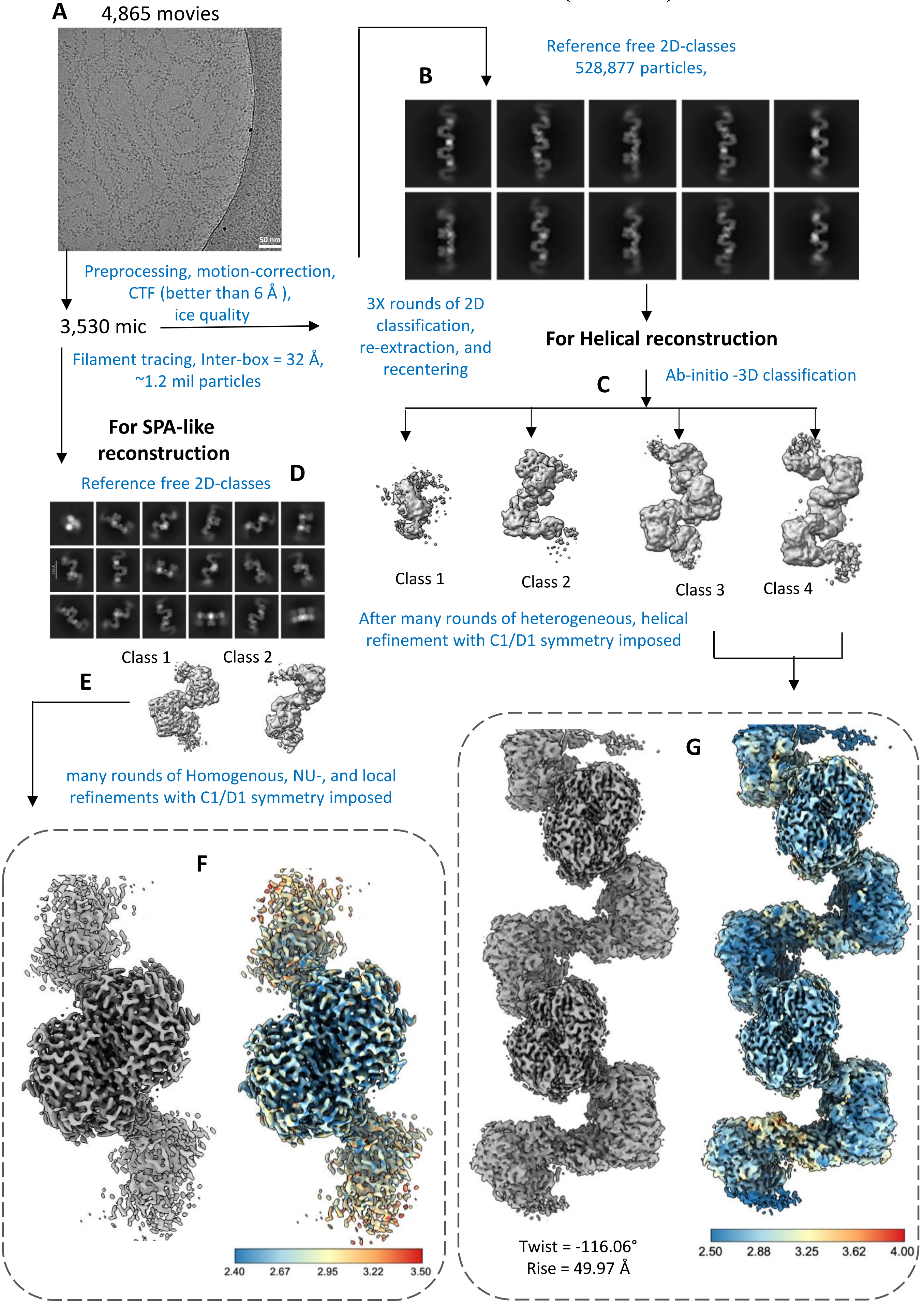


Supplementary figure 1

### The *trans*-basal CBS filament (dataset 1)

#### Helical reconstruction

**H**

D1 symmetry imposed

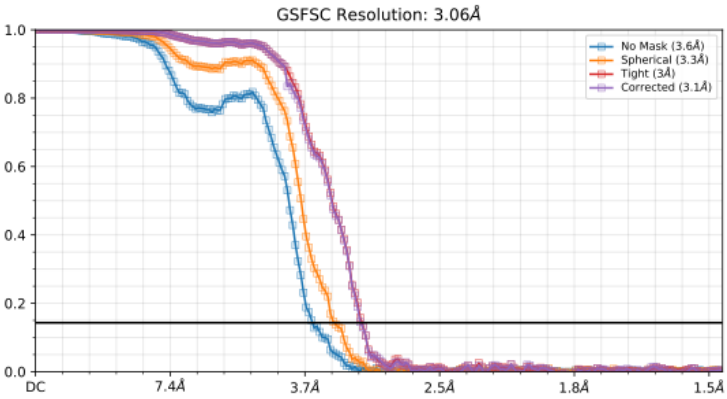


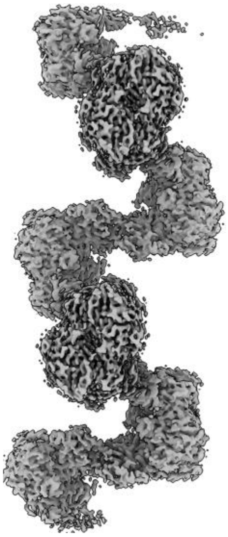

Twist = -116.06°
Rise = 49.97 Å

#### SPA-like reconstruction

**I**

D1 symmetry imposed

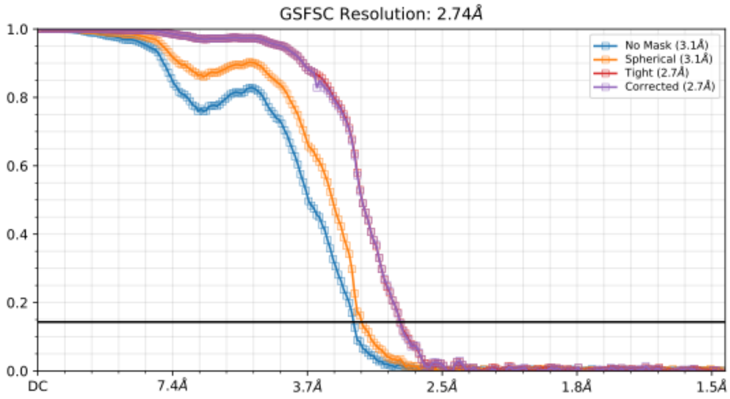


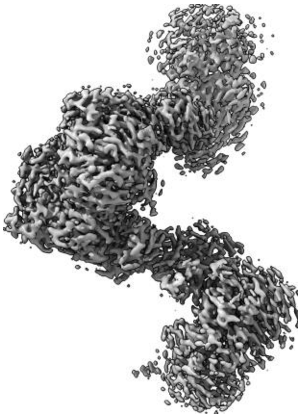

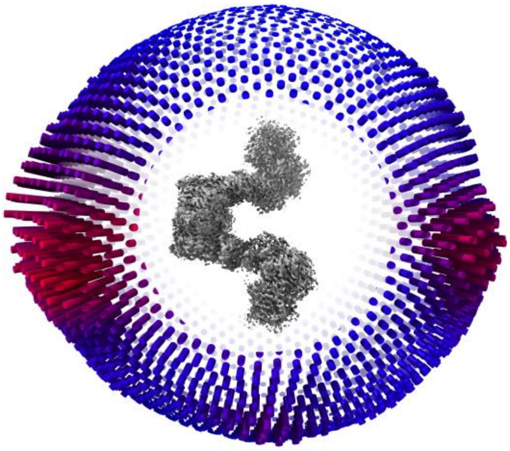

**Supplementary figure 1 (continued)**

**Supplementary figure 2. Cryo-EM analysis for determination of helical parameters of the *trans*-basal CBS filament. A** – Ab initio helical reconstruction of a CBS filament containing approximately 14–16 dimeric repeating units, used to estimate the helical pitch and rise in real space with UCSF Chimera. **B** – Representative 2D class average of the same long filament together with its corresponding power spectrum, used to calculate the helical parameters in the Fourier domain. Both real-space and Fourier analyses provided consistent initial estimates of ~164 Å pitch and ~54 Å rise. **C** – Validation of these parameters by refining and reconstructing a longer filament comprising 12–14 dimer repeats. The resulting map, fitted model, and FSC curve are shown, reaching a resolution of 4.71 Å.

### The *trans*-basal CBS filament (dataset 1)

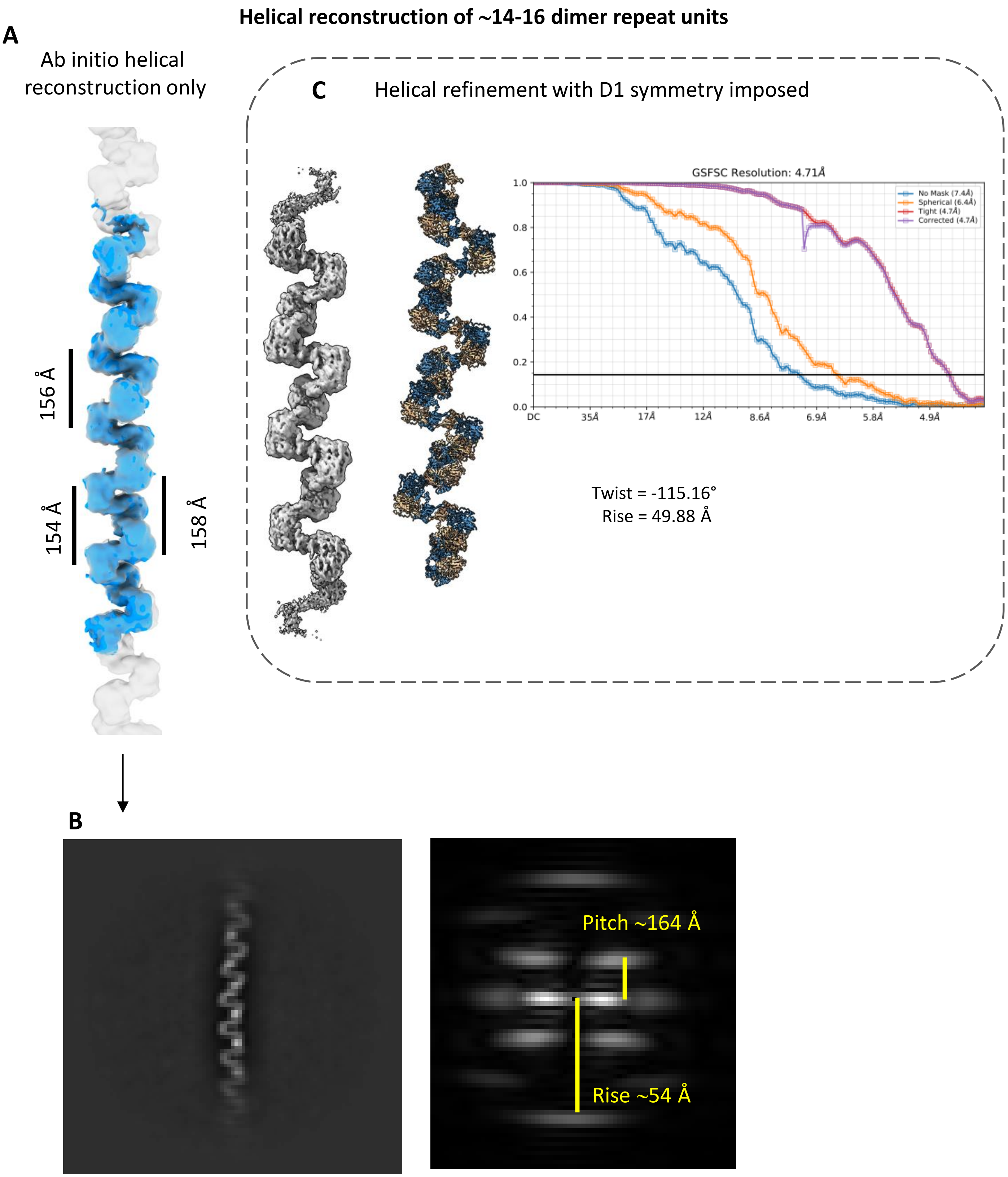


**Supplementary figure 2**

**Supplementary figure 3. Cryo-EM image processing workflow for structural elucidation of human CBS in the presence of its substrate L-serine.** **A** – Processing pipeline combining both helical and single-particle analysis (SPA)-like approaches. Representative reference-free 2D class averages are shown. Particles contributing to a consensus map were subsequently used to sort and classify substates by both strategies: SPA classification with a focused mask around one CD dimer repeat and four RDs, and helical classification with a mask covering three dimer repeats. **B&C** – This approach resolved three well-defined structural classes, corresponding to the key catalytic intermediates: CBS-Lys-PLP (class 3), CBS-PLP-Ser (class 4), and CBS-PLP-AA (class 5).

### The Ser-*trans*-basal CBS filament (dataset 2)

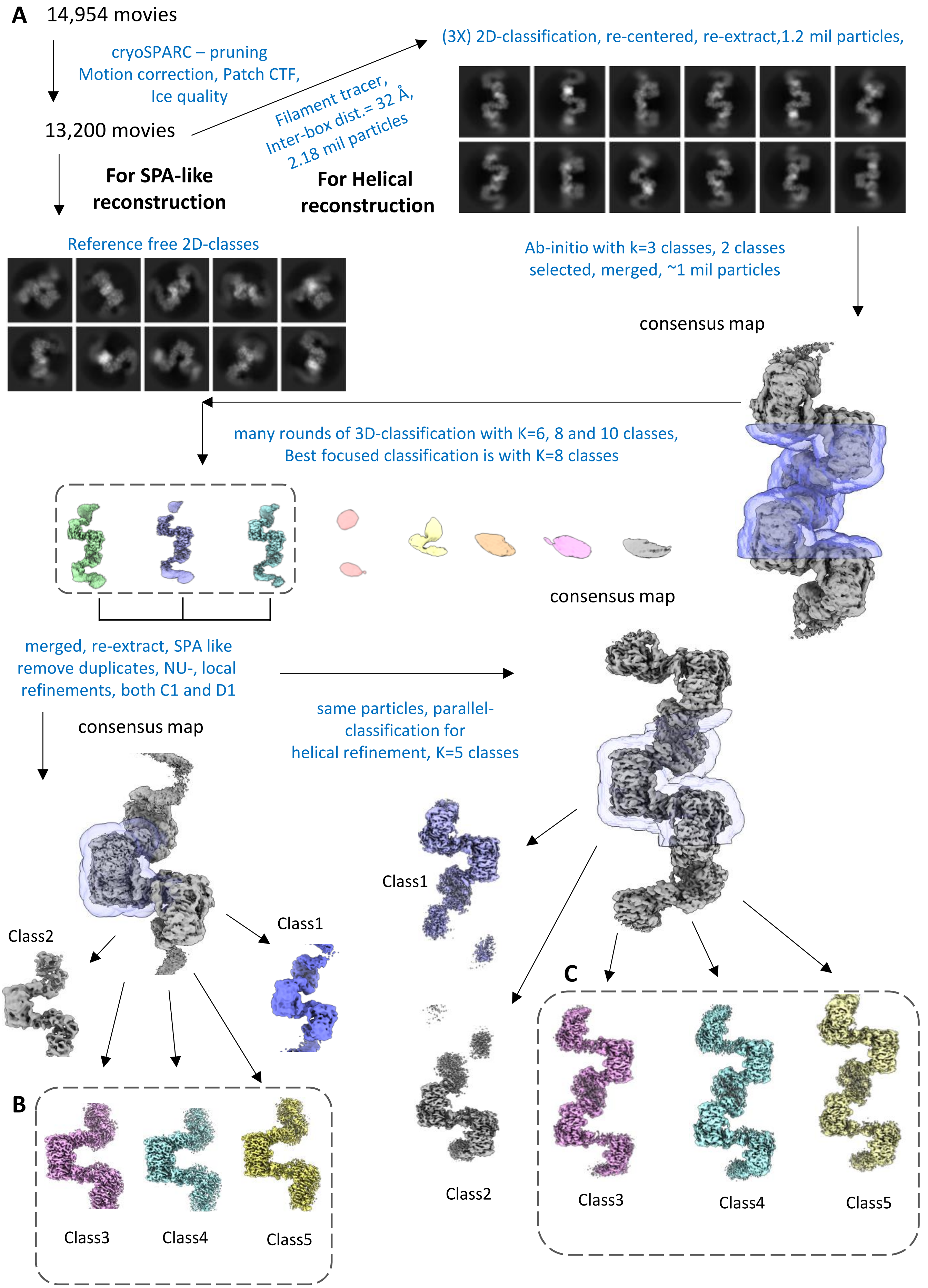


Supplementary figure 3

**Supplementary figure 4. Cryo-EM helical reconstructions of L-serine substrate-bound CBS *trans*-basal filaments.** **A** – CBS-PLP-AA (class 5), representing the PLP–aminoacrylate intermediate, resolved at 2.62 Å. The panel includes the local resolution map and corresponding FSC curve. **B** – CBS-PLP-Ser (class 4), representing the PLP–serine external aldimine intermediate, resolved at 2.92 Å. The global resolution is shown by the FSC curve alongside the local resolution map.

### The Ser-*trans*-basal CBS filament (dataset 2)

#### Helical reconstruction

**A**

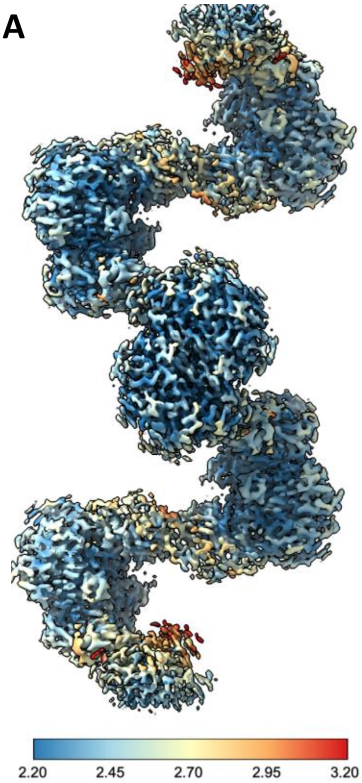


Class5, D1 symmetry imposed,
Twist = -115.312°, Rise = 49.940 Å

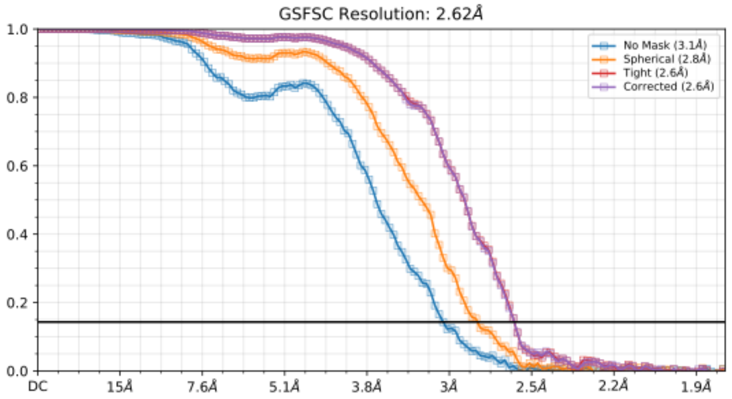


**B**

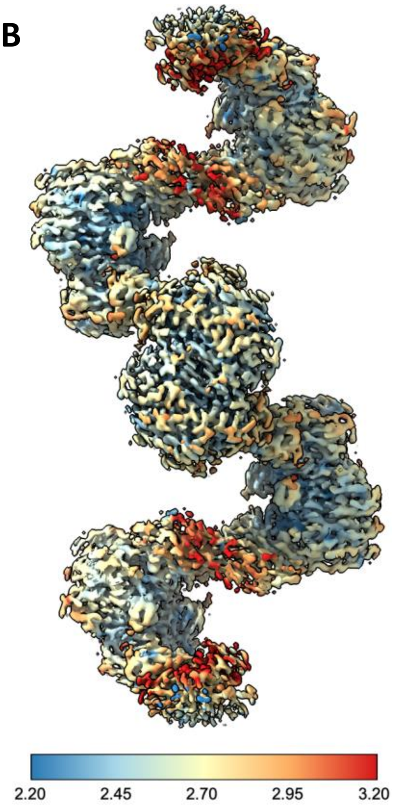


Class4, D1 symmetry imposed,
Twist = -114.984°, Rise = 50.78 Å

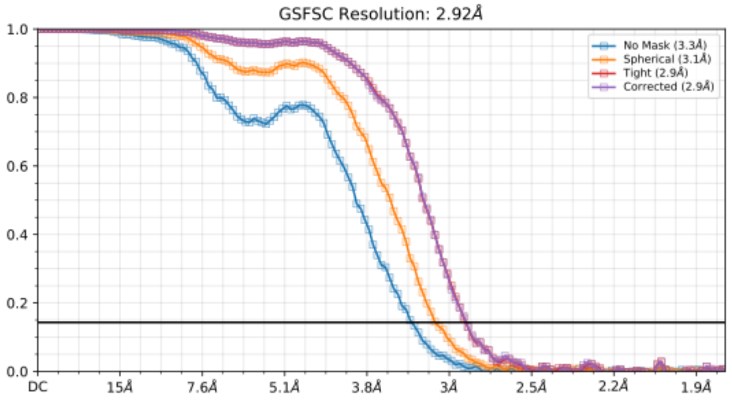


**Supplementary figure 4**

**Supplementary figure 5. Cryo-EM single-particle reconstructions of L-serine substrate-bound CBS *trans*-basal filaments.** **A** – CBS-PLP-AA (class 5), corresponding to the PLP–aminoacrylate intermediate, reconstructed at 2.08 Å resolution. The panel shows the local resolution map, FSC curve, and particle orientation distribution. **B** – CBS-PLP-Ser (class 4), corresponding to the PLP–serine external aldimine intermediate, reconstructed at 2.20 Å resolution. The global resolution is validated by the FSC curve, with accompanying local resolution and particle orientation distribution maps.

### The Ser-*trans*-basal CBS filament (dataset 2)

#### SPA-like reconstruction

A

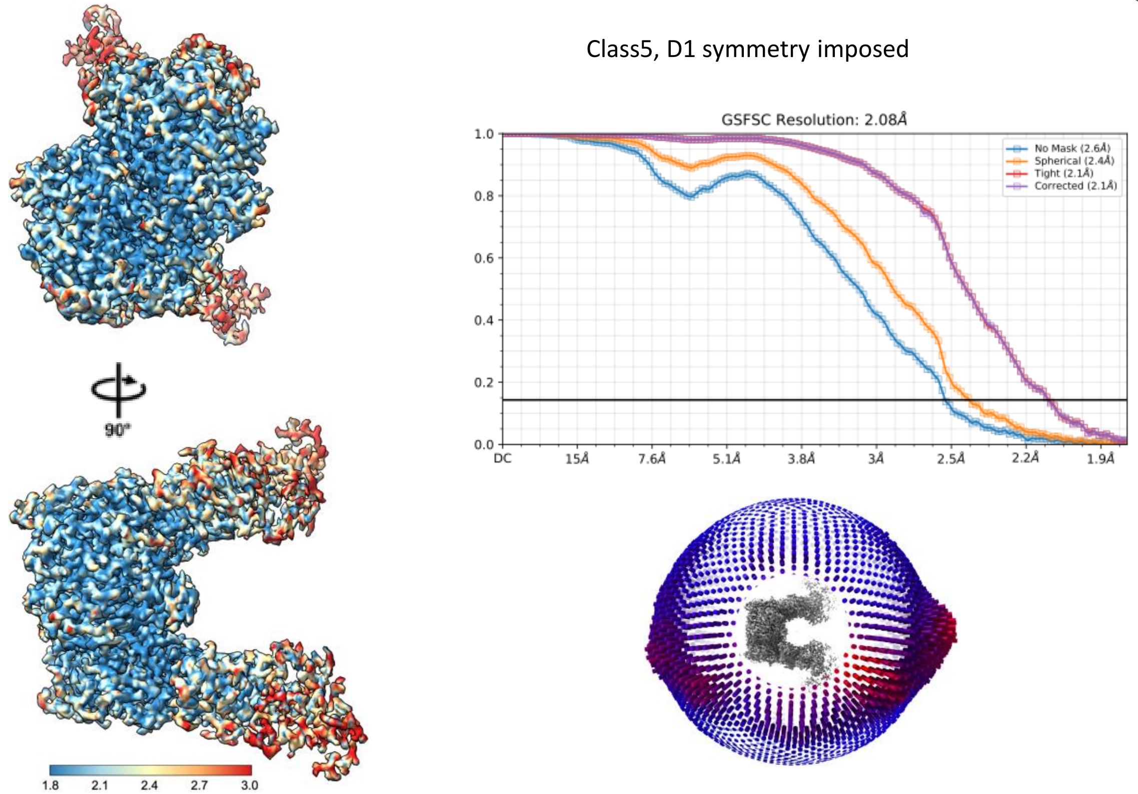


B

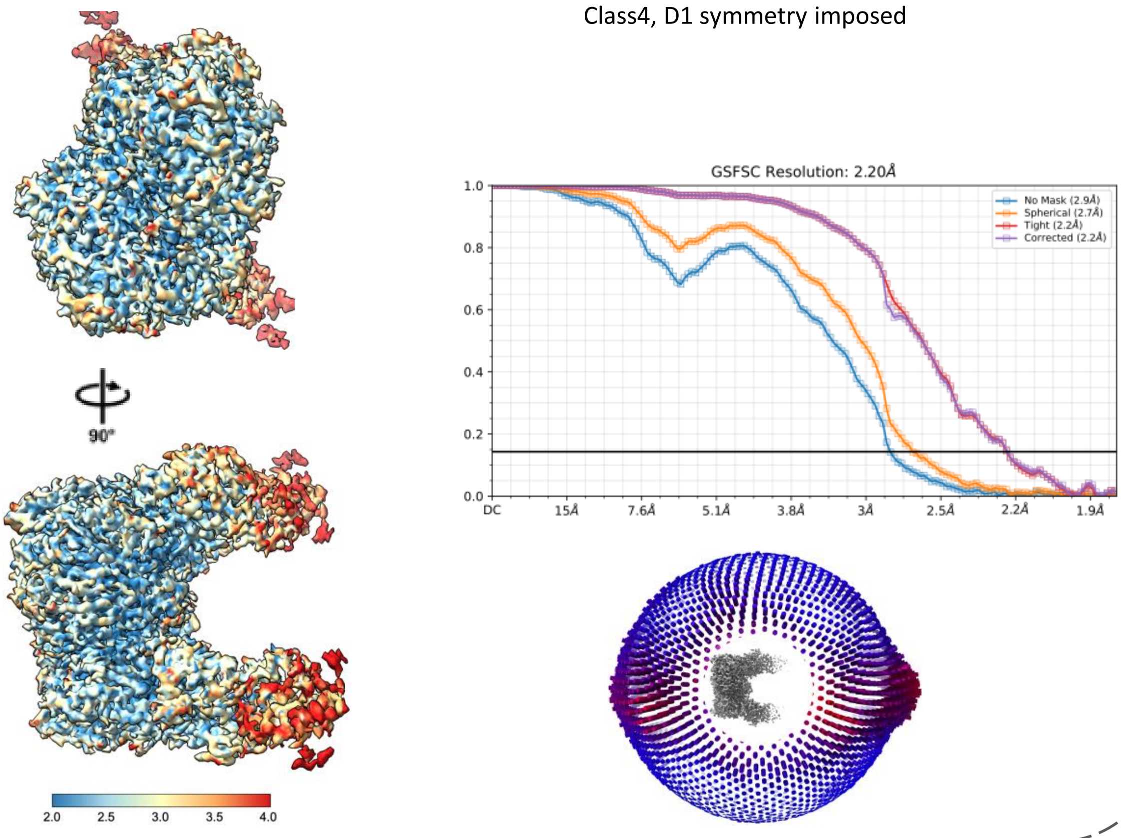


Supplementary figure 5

**Supplementary figure 6. Cryo-EM image processing workflow of the *cis*-basal CBS filament in the presence of a non-activating allosteric ligand sinefungin/adenosylornithine (SAO). A** – Representative electron micrograph of human CBS incubated with 0.5 mM SAO, showing long filamentous assemblies with a distinct morphology compared to the *trans*-basal state. This is further illustrated in the corresponding 2D class averages. The processing workflow combined both helical reconstruction and SPA-like approaches, beginning with reference-free 2D class averaging. Particles contributing to a consensus reconstruction were subsequently sorted and classified to exclude suboptimal images. **B&C** – Helical classification and refinement strategies. Panel **B** shows refinement using masks covering the entire repeats of both CDs and RDs, while panel **C** focuses on refinement with a localized mask around the central RDs only.

### The SAO-bound *cis*-basal CBS filament (dataset 3)

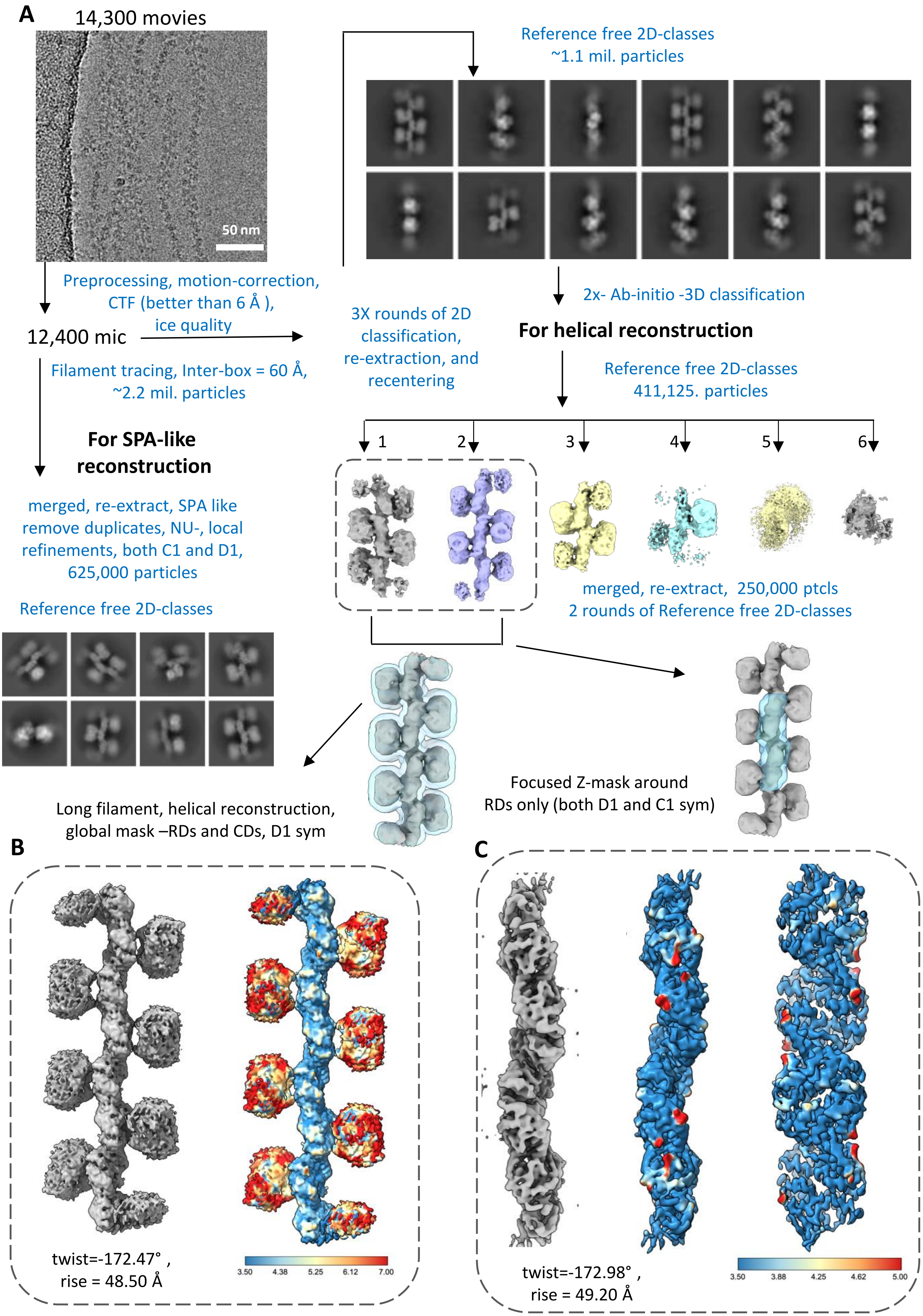


Supplementary figure 6

**Supplementary figure 7. Cryo-EM helical reconstructions of SAO-bound *cis*-basal CBS filament. A** – Helical refinement using masks covering the entire repeats of both CDs and RDs, yielding a reconstruction at 4.86 Å resolution as indicated by the FSC curve. **B** – Focused refinement with a localized Z-mask applied around the central RDs, which improved the map quality and allowed clearer visualization of the bound SAO ligand. The corresponding FSC curve is shown.

### The SAO-bound *cis*-basal CBS filament (dataset 3)

#### Helical reconstruction

**A**

Long filament, helical reconstruction, global mask –RDs and CDs, D1 sym

Twist = -172.47°
Rise = 48.50 Å

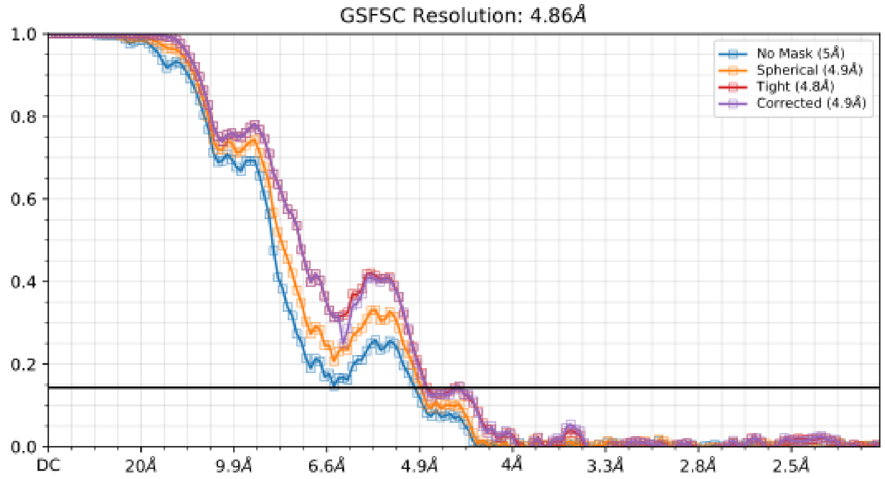


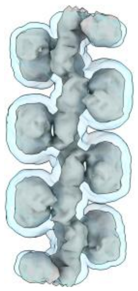

**B**

Focused Z-mask around RDs only (both D1- and C2-sym)

Twist = -172.98°
Rise = 49.20 Å

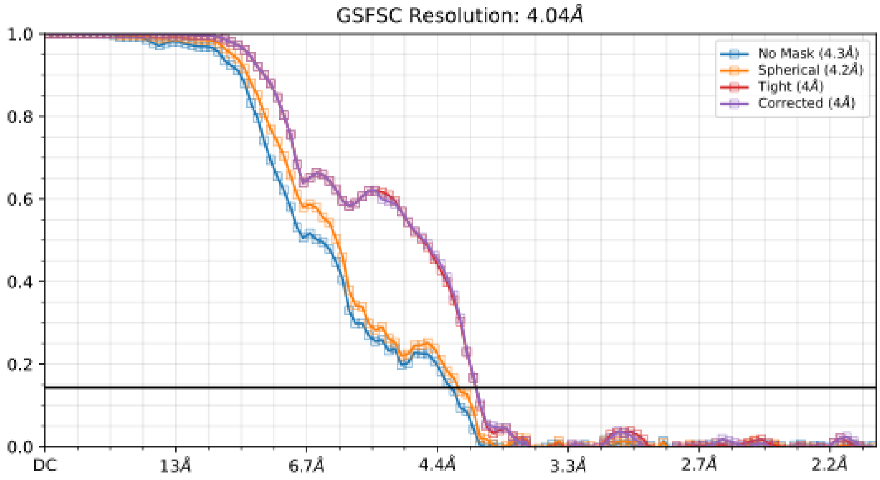


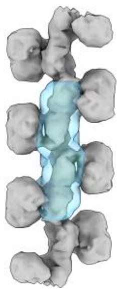

Supplementary figure 7

**Supplementary figure 8. Cryo-EM single-particle reconstruction of cis-basal CBS in the presence of the SAO ligand. A&B** – Representative 2D class averages (**A**) and the resulting 3D reconstruction (**B**) obtained using a focused mask encompassing one CD and six RD domains. The local resolution map, refined to 4.1 Å, shows that the CD is resolved at slightly lower resolution compared to the central RD region. **C** –FSC curve and particle orientation distribution corresponding to the reconstruction.

### The SAO-bound *cis*-basal CBS filament (dataset 3)

**SPA-like reconstruction**

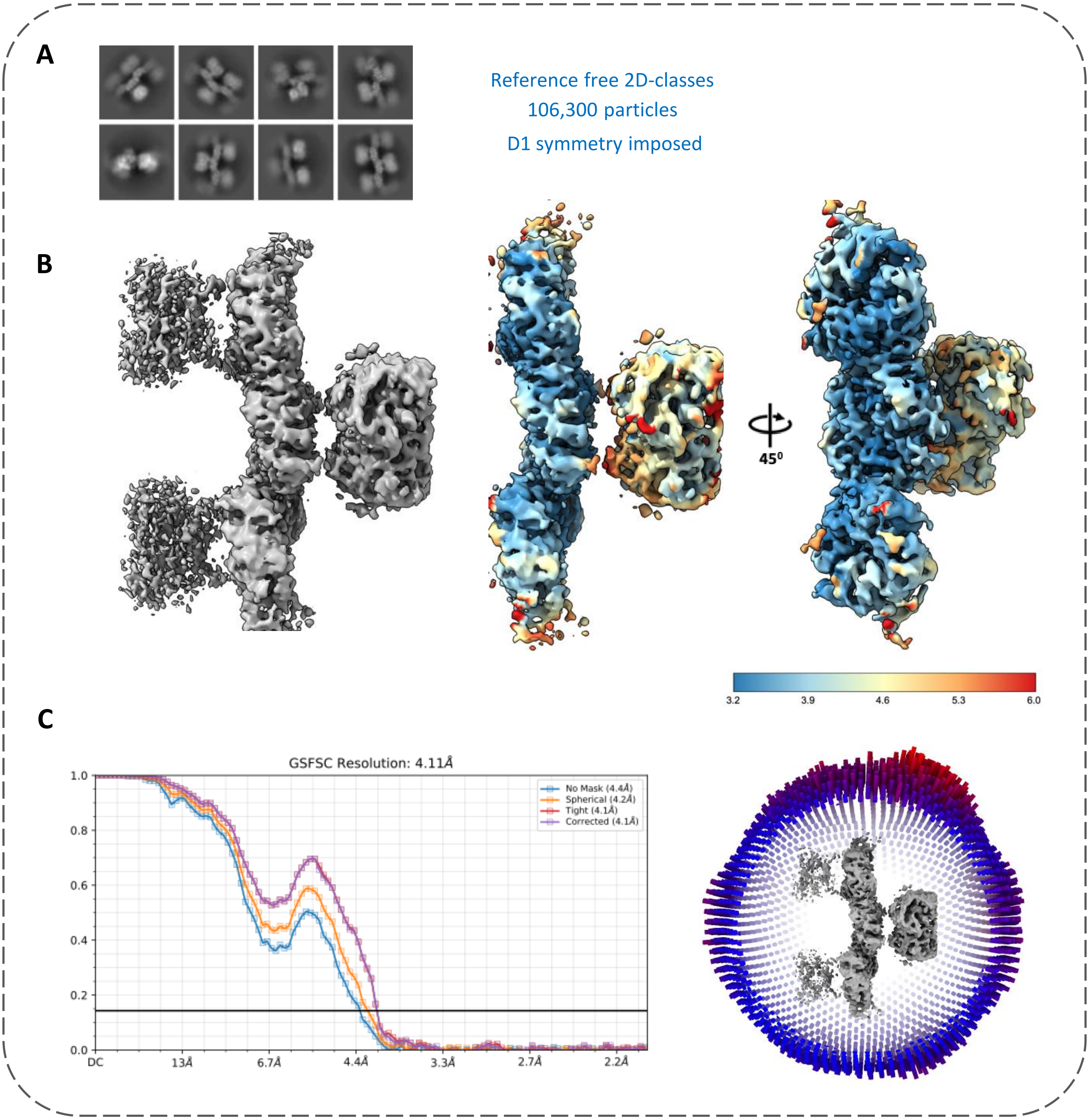


Supplementary figure 8

**Supplementary figure 9. Cryo-EM image processing workflow for structural elucidation of human CBS in the presence of an activating allosteric ligand S-adenosylmethionine (SAM). A** – Representative electron micrograph showing higher-order filamentous assemblies of human full-length CBS in the presence of 0.5 mM SAM (highlighted with yellow markers). These assemblies display distinct morphology compared to both the *trans*-basal and SAO-bound *cis*-basal CBS filaments. **B&C** – Representative 2D class averages (**B**) and 3D classification density maps (**C**) obtained from ab initio and helical reconstructions performed without imposing helical parameters. **D** – After multiple rounds of 3D classification, the best classes were merged, re-extracted with a larger box size, and subjected to additional 2D classification to exclude suboptimal particles. The resulting averages reveal stacked filaments with a core formed uniquely rearranged CDs termed "*allo*-CDs" (yellow arrows).

### The SAM-bound *allo*-activated stacked CBS filament (major fraction; dataset 4)

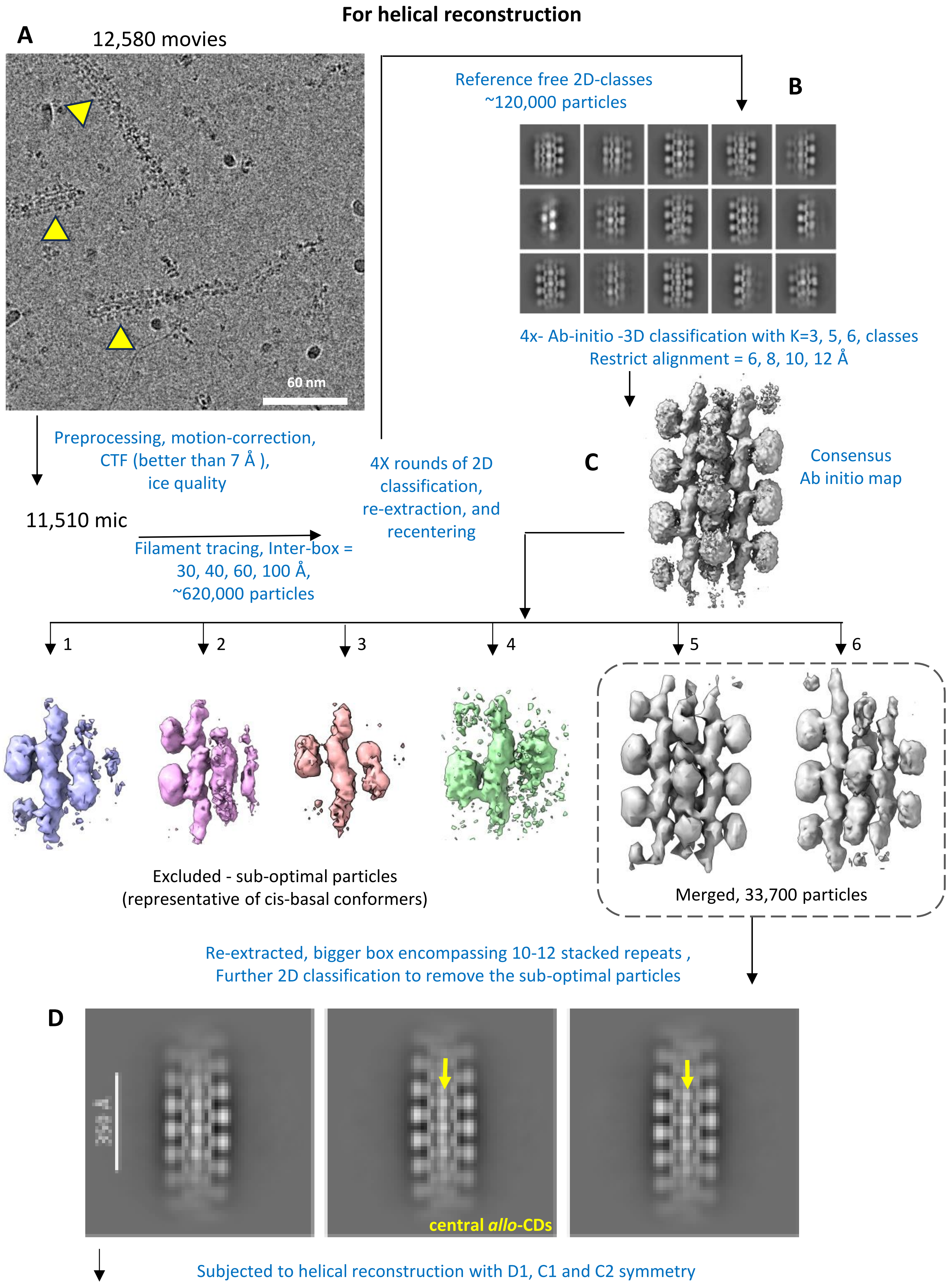


Supplementary figure 9

**Supplementary figure 10. Cryo-EM helical reconstructions of the SAM-bound *allo*-activated stacked CBS filament.** **A** – Helical reconstruction and refinement performed using masks covering two consecutive repeats of both CDs and RDs. Reconstruction was carried out without imposing symmetry (C1) to allow visualization of the central *allo*-CD region, which is stacked between the two incoming filaments. The CD-RD connecting linker motifs, RDs, outer CDs, and central *allo*-CDs are designated. **B** – The final reconstruction was resolved at 8.43 Å, as indicated by the FSC curve. The corresponding local resolution map shows that the flanking RDs are better defined compared to the central *allo*-CDs.

### The SAM-bound *allo*-activated stacked CBS filament (major fraction; dataset 4)

#### Helical reconstruction

Final helical reconstruction = after many rounds of D1 symmetry and followed by C1 symmetry

**A**

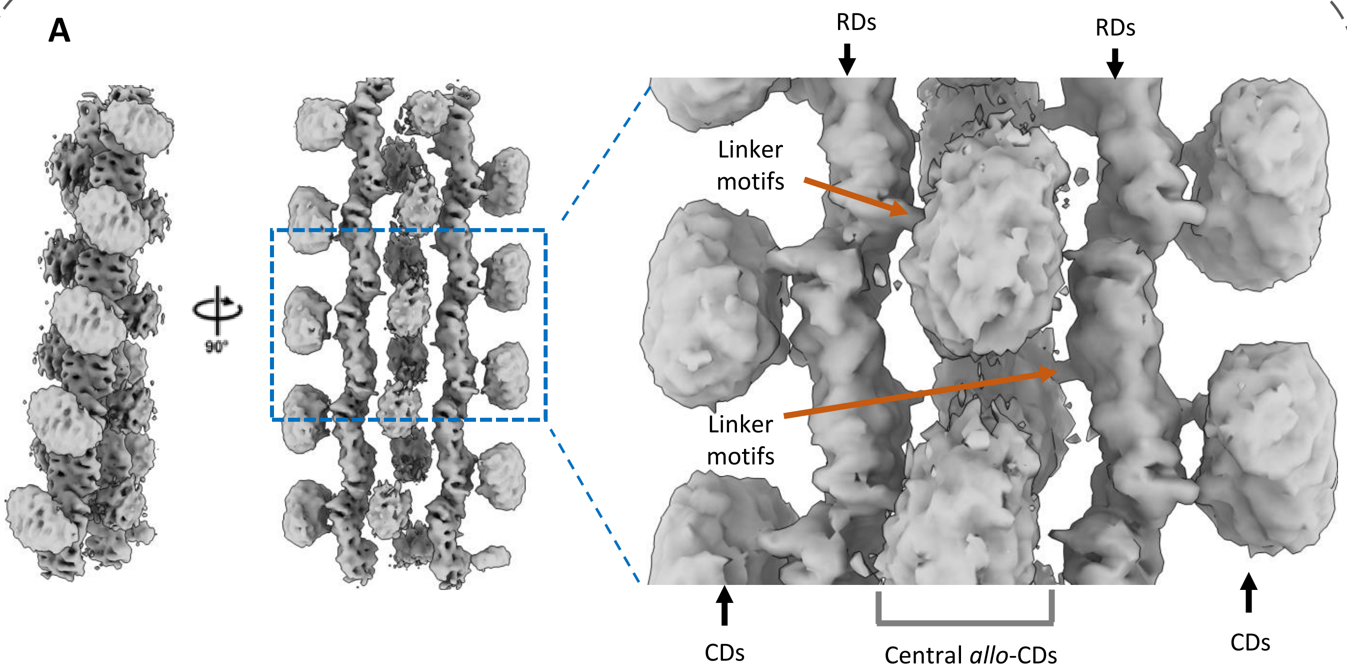


C1 symmetry imposed

**B**

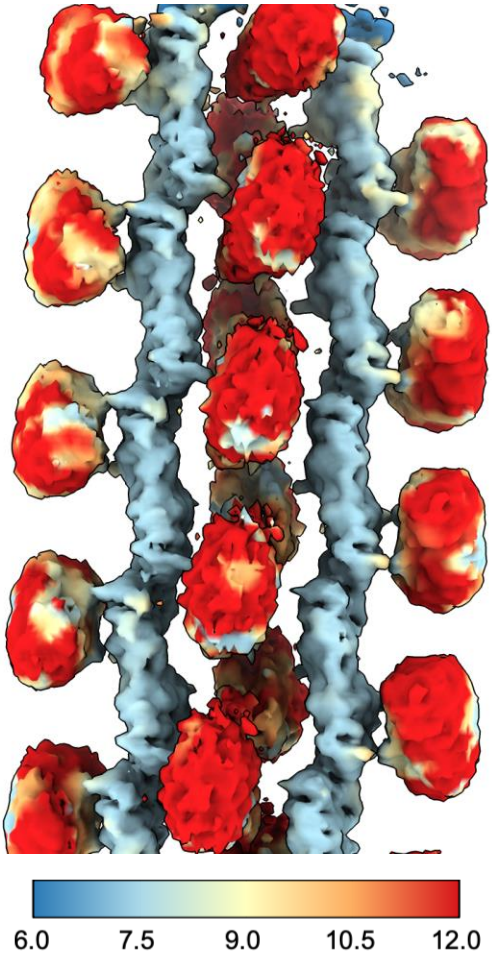


Twist = -176.08°
Rise = 48.28 Å

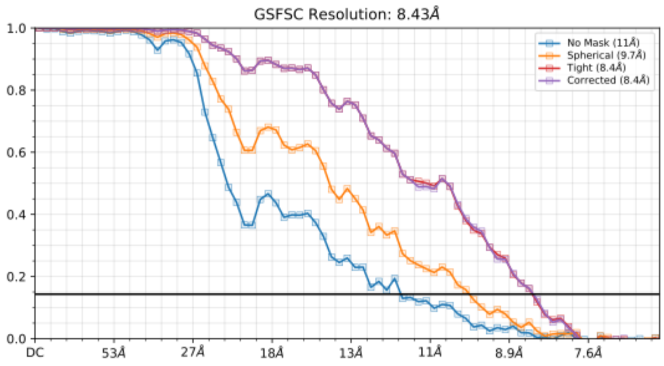


Supplementary figure 10

**Supplementary Fig. 11. Complementary cryo-EM helical reconstruction of the SAM-bound *allo*-activated stacked CBS filament with C2 symmetry imposed.** **A** – Helical reconstruction and refinement carried out with C2 symmetry applied, aimed at improving the resolution of the central *allo*-CD region stacked between the two incoming filaments. The RDs, CDs, and central *allo*-CDs are highlighted. **B** – The reconstruction reached 9.65 Å resolution, as indicated by the FSC curve. The corresponding local resolution map shows no improvement in density for the central *allo*-CDs compared to the C1 reconstruction (Supplementary figure 10), underscoring its intrinsic flexibility.

### The SAM-bound *allo*-activated stacked CBS filament (major fraction; dataset 4)

#### Helical reconstruction

**A** Helical reconstruction = C2 symmetry imposed to resolve the central CDs

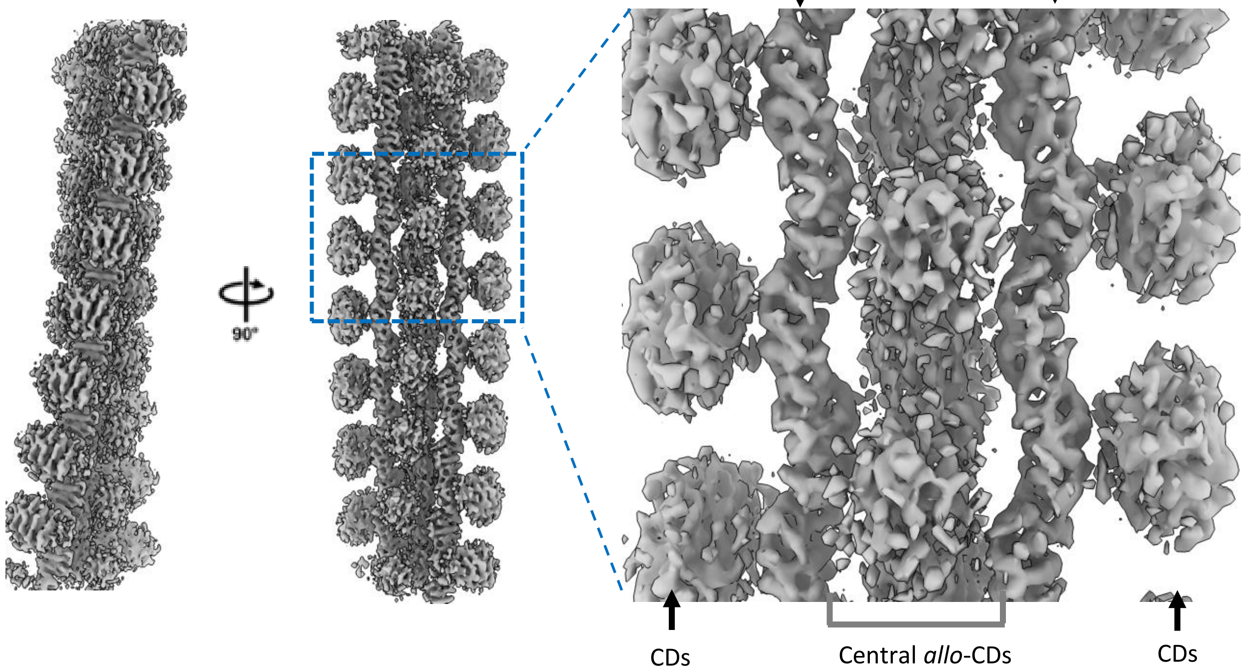


**B** C2 symmetry imposed

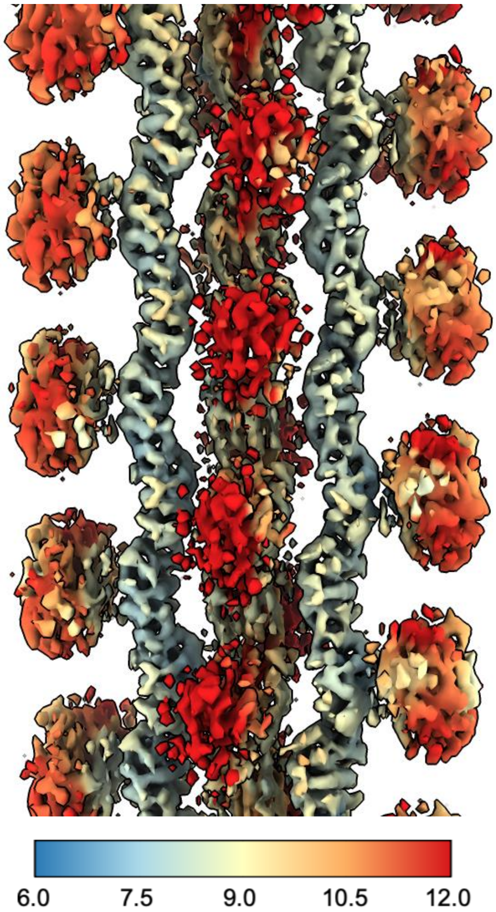


Twist = -174.766°
Rise = 48.150 Å

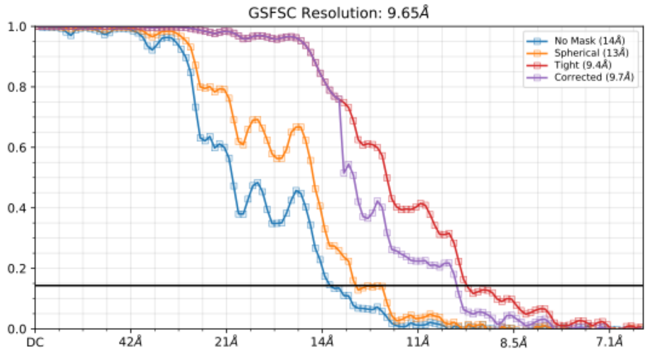

**Supplementary figure 12. Cryo-EM single-particle reconstruction of the SAM-bound *allo*-activated stacked CBS filament.** **A** – Representative 2D class averages showing end-on and tilted views of the SAM-bound *allo*-activated stacked CBS filament. Yellow arrows highlight the central *allo*-CDs. **B** – Focused refinement using a localized Z-mask around the central *allo*-CD region with C2 symmetry imposed. **C** – Subsequent refinement with a full mask under C1 symmetry improved overall map quality, enabling clearer visualization of the central *allo*-CD stalk. The corresponding FSC curve and local resolution map are shown. This strategy yielded a global reconstruction at 7.87 Å resolution.

### The SAM-bound *allo*-activated stacked CBS filament (major fraction; dataset 4)

**SPA-like reconstruction**

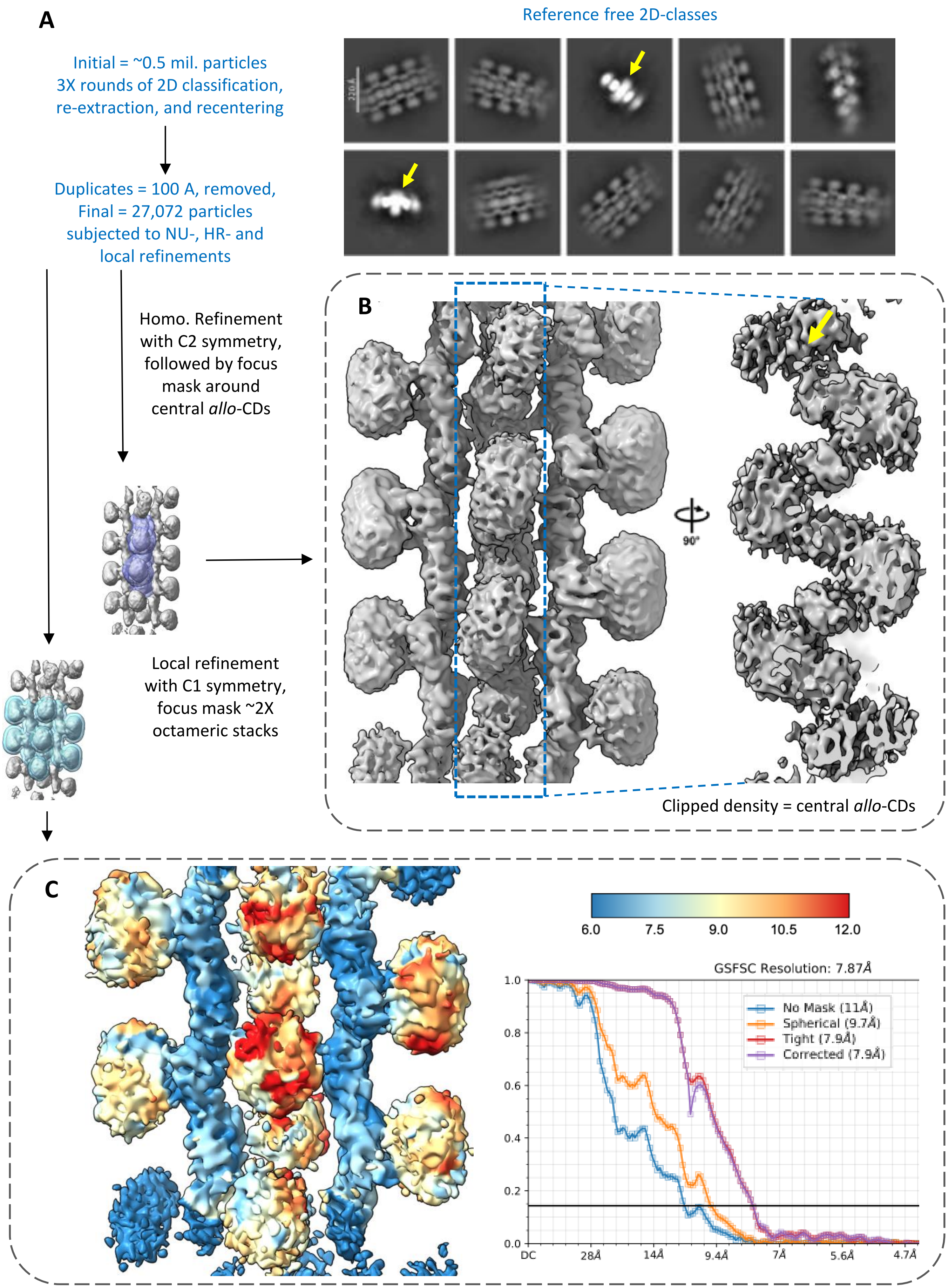


**Supplementary figure 12**

**Supplementary figure 13. Cryo-EM reconstructions of cis-basal CBS in the presence of SAM. A** – Helical reconstruction approach revealing a minor population of SAM-bound *cis*-basal CBS filaments. Representative 2D class averages, the corresponding 3D reconstruction, local resolution map, and FSC curve are shown, yielding a final resolution of 6.76 Å. **B** – SPA-like reconstruction of the SAM-bound *cis*-basal CBS filament using a focused mask around the central RD region. The corresponding FSC plots and local resolution maps are shown, demonstrating that the RDs are better resolved than the CDs. This refinement yielded a map at 4.12 Å resolution.

### The SAM-bound *cis*-basal CBS filament (minor fraction; dataset 4)

#### Helical reconstruction

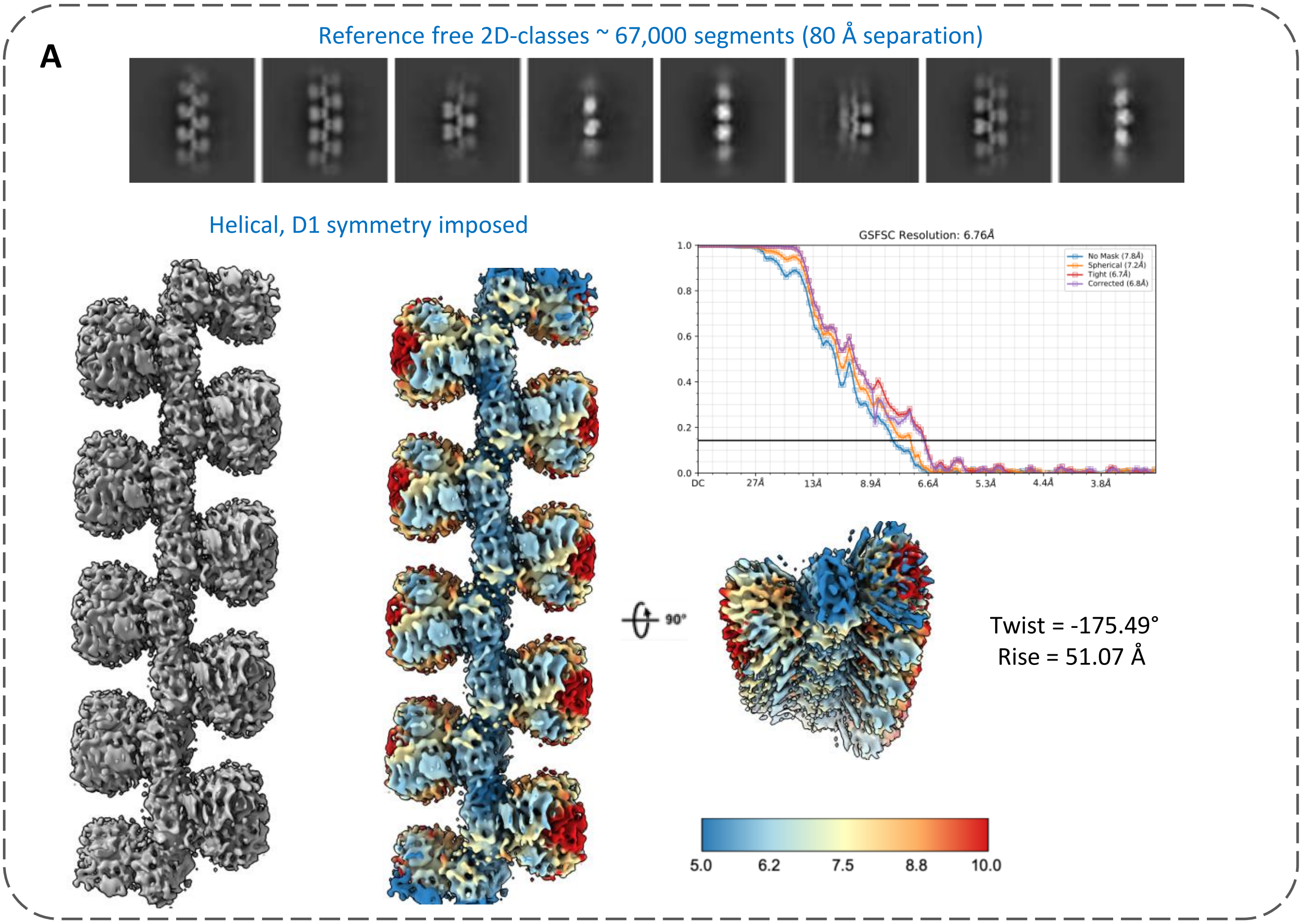


#### SPA-like reconstruction

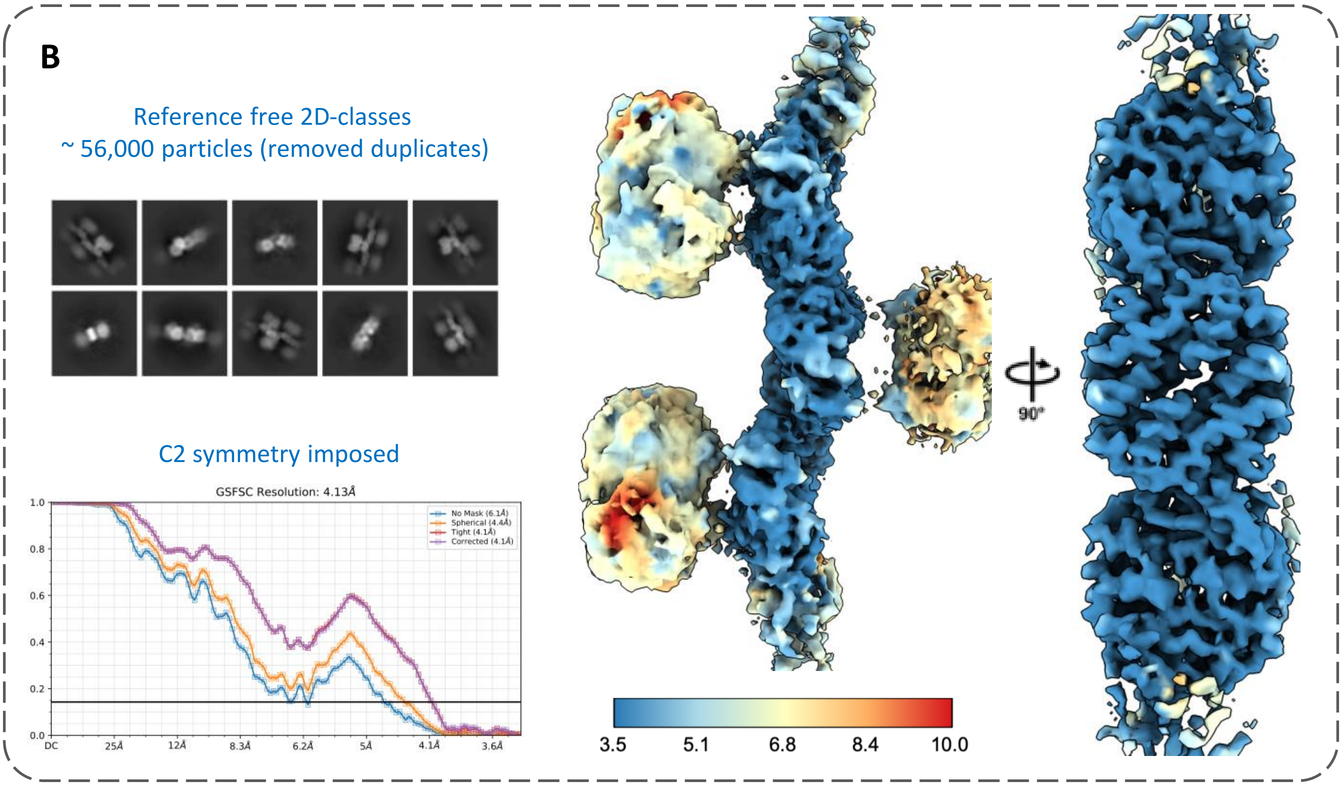


Supplementary figure 13

**Supplementary figure 14. Additional molecular assemblies and disassembly states observed in the SAO and SAM datasets.** **A** – Broken dimeric assemblies containing two CDs and two RDs identified in both the SAO- and SAM-bound datasets. Below, extended dimers consisting of two CDs and four RDs were also observed. **B** – A unique assembly observed only in the SAO dataset, possibly representing a transitional intermediate captured during the shift from the ligand-free *trans*-basal to the SAO-bound *cis*-basal state. **C** – Long filamentous assemblies of RDs detected in 2D class averages. **D** – Isolated densities corresponding to broken central *allo*-CDs observed only in the SAM-bound dataset, likely representing disassembled intermediates of the resolved *allo*-activated stacked CBS filament.

### Other molecular CBS species observed in of SAO- and SAM-bound datasets

**A**

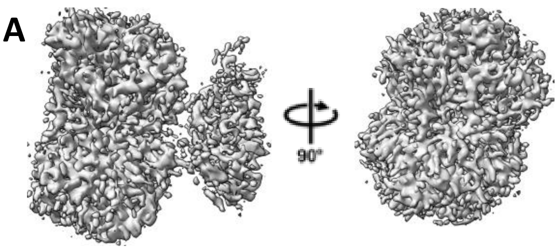


Separated *cis*-basal dimer
(observed in both the SAO- and SAM-bound datasets)

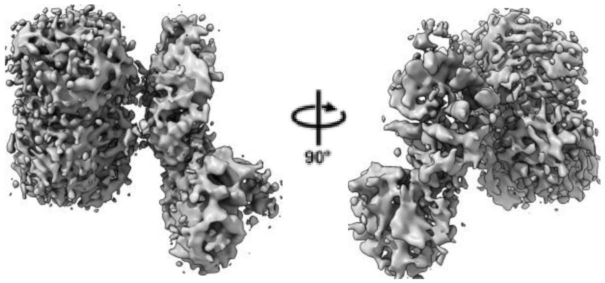


Separated *cis*-basal dimer + 2 RDs
(observed in both the SAO- and SAM-bound datasets)

**B**

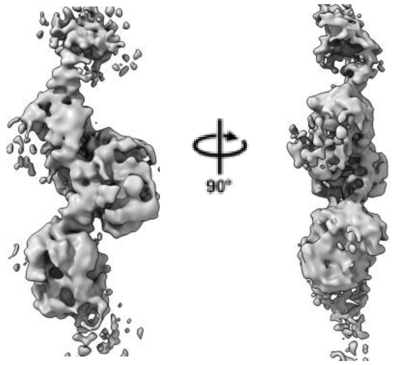


Transition from *trans*-basal to SAO-bound *cis*-basal CBS filament (observed in SAO-bound dataset)

**C**

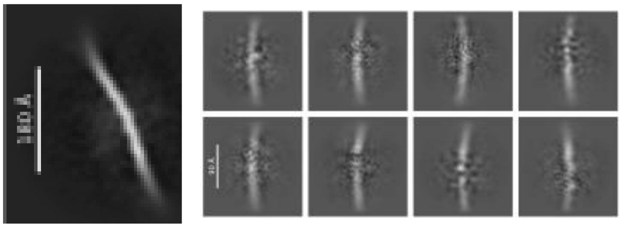

Broken RDs
(observed mostly in SAM-bound dataset)

**D**

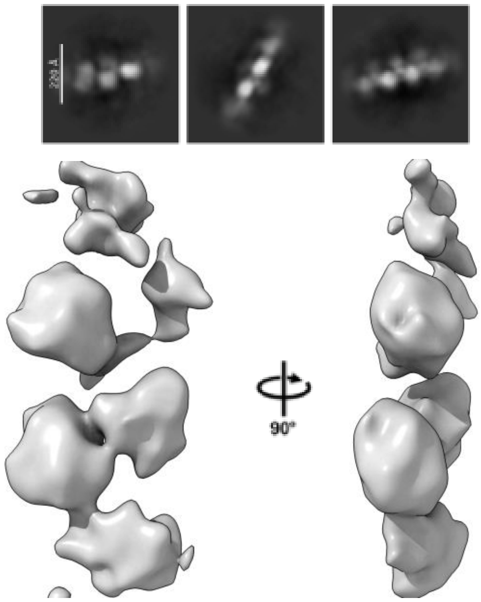


Broken central *allo*-CDs
(observed only in SAM-bound dataset)

**Supplementary figure 14**
